# Ribosome profiling reveals translational dynamics between an archaeal virus and its host

**DOI:** 10.64898/2026.09.13.751334

**Authors:** Matthew F. Isada, Pavlína Gregorová, Palak Sadana, L. Peter Sarin, Jocelyne DiRuggiero

**Affiliations:** Johns Hopkins University; University of Helsinki

**Keywords:** Archaea, translation, ribosome profiling, host-virus interactions, tRNA

## Abstract

Host-virus interactions remain poorly understood in archaea, whose information processing systems harbor a mosaic of bacterial and eukaryotic features. Here, we used complementary multiomics to investigate interactions between *Haloferax* tailed virus 1 (HFTV1) and the halophilic archaeon *Haloferax gibbonsii* (*Hg*), capturing global translational and transcriptional dynamics across the infection cycle. This revealed co-regulatory expression patterns across HFTV1 and *Hg* genes. Post-transcriptional regulation was observed for several early HFTV1 genes and the late HFTV1 gene *gp17*, and the HFTV1 tRNA co-sedimented with *Hg* ribosome complexes. Infection reshaped the translational landscape of *Hg*, causing a slow decline in information processing activity, a complex transcriptional response, and amino acid starvation. An unannotated Tat-dependent fimbrial system of *Hg* was co-regulated with early HFTV1 genes and was correlated with increased biofilm formation, suggesting a potential antiviral defense system. These findings provide high-resolution insight into archaeal host-virus interactions at the evolutionary interface between bacteria and eukaryotes.

## INTRODUCTION

Organisms across the Tree of Life are subject to viral predation, which ultimately regulates community dynamics, drives evolution, and shapes the global ecosystem [1]. The infection cycle of each virus is mediated by molecular interactions between components of host and viral origin. However, while the molecular host-virus interactions have been extensively characterized in bacteria and eukaryotes, they remain understudied in archaea, an environmentally and evolutionarily significant form of life. Although archaea famously dominate extreme environments, they are also abundant in soil [2], fresh water [3], and marine ecosystems, performing crucial steps in biogeochemical cycling [4–6]. Archaea share a close evolutionary relationship with eukaryotes, underscored by the discovery of Asgard archaea and their potential role in eukaryogenesis [7, 8]. This relationship is supported by the mosaic of eukaryotic and prokaryotic features within the archaeal information processing system: archaea possess eukaryotic homologs of replication, transcription, and translation machinery [9–11], yet their transcripts are uncapped [11], sometimes polycistronic [12, 13] or leaderless [13, 14], and expressed by coupled transcription-translation machinery like bacteria [11]. Consequently, archaeal viruses must encounter features reminiscent of both eukaryotes and bacteria in their hosts, potentially resulting in an evolutionary mosaic of host-virus interactions.

Archaeal viruses are incredibly diverse. Some have virion morphologies [15, 16] and assembly mechanisms [17–23] that are entirely unique to archaea. In contrast, tailed and untailed icosahedral virions are similar in structure, assembly, and genome across archaeal, bacterial, and eukaryotic hosts [15, 16, 24–28], hinting at the descendance of all icosahedral viruses from a common ancestor [16, 29, 30]. In egress, virions can bud from the cell membrane like eukaryotic viruses [23, 31], virions can be released by large pyramidal structures that transect the membrane and S-layer [32–34], or the cell membrane can be disrupted by unknown mechanisms [35]. Even within Asgard archaea, the ancestors of eukaryotes [7, 8], the metagenomic signals of their associated viruses vary dramatically: some are similar to phages [36] or eukaryotic viruses [37], while others are specific to archaea [36]. This wealth of information comes from extensive research using genomic, structural, and microscopy-based techniques. However, many isolated host-virus systems in archaea suffer from limited tractability, requiring extreme growth conditions, following noncanonical infection cycles, or lacking genetic and molecular research tools. This has hindered the study of several aspects of archaeal host-virus interactions, including host and viral gene expression during infection. Only a handful of studies have applied transcriptomics to comprehensively study gene expression in archaeal host-virus systems. These studies showed that archaeal viruses have varying effects on host defense systems: the lytic *Sulfolobus* virus SIRV2 [38], the lytic *Methanosarcina* virus MetSV [39], and the "carrier state" *Sulfolobus* virus SIRV3 induce CRISPR-Cas systems [38, 40], toxin-antitoxin systems [38], and/or restriction-modification systems [39], whereas the chronic virus LSV-48N inhibits CRISPR spacer acquisition [41] and the chronic virus HFPV-1 down-regulates the expression of host proviruses [42]. Additionally, infection by the lytic virus MetSV increases expression of host proteins involved in DNA replication and protein folding [39]. These findings are consistent with canonical activities of viral infection known from bacteria and eukaryotes, such as induction or suppression of host defense systems and the redirection of host resources to facilitate viral genome and protein synthesis.

These few transcriptomic studies have provided valuable insight into archaeal host-virus interactions. However, some of the studies were performed at low temporal resolution, with only 2-3 time points collected during infection [39, 41, 42]. Additionally, many genes from both the host and viral genomes remain uncharacterized, accounting on average for 40% of any given archaeal genome [43] and 40-80% of the viral genomes in the transcriptomic studies listed above. Since many genes will inevitably be uncharacterized in these systems, identifying ones that may mediate host-virus interactions could be improved by first searching for signs of co-regulation to select genes for further investigation. Transcriptomic studies have also how viral gene expression is regulated in archaea. Some level of regulation must inevitably occur at the transcriptional level; unlike many bacterial and eukaryotic viruses, no known archaeal viruses encode an RNA polymerase (RNAP), making viral transcription in archaea heavily reliant on host machinery [44]. Redirection of host transcription may be facilitated by the presence of host-like promoter elements in the genomes of archaeal viruses [44–46]. Additionally, biochemical studies showed that a few viral proteins are involved in transcriptional regulation [47]: the *Acidianus* virus ATV encodes a global transcriptional repressor that associates directly with the host RNAP [44] while the *Acidianus* virus AFV6 [48], the *Sulfolobus* virus SIRV1 [49, 50], and the *Sulfolobus* virus SSV1 [51, 52] each encode gene-specific transcriptional activators or repressors. However, most targets of the gene-specific regulators are either uncharacterized or were determined *in vitro*, leaving their roles in infection uncertain. Furthermore, nothing is known for archaeal viruses about post-transcriptional regulation, a process that can strongly influence viral gene expression in bacteria [53–55] and eukaryotes [56].

One promising system to capture multiple levels of gene expression during the infection cycle is the lytic *Haloferax* tailed virus 1 (HFTV1) and its host, the halophilic archaeon *Haloferax gibbonsii* LR2-5 (*Hg*). This is a highly tractable system, with *Hg* closely related to the established model archaeon *Haloferax volcanii* [57]. *Hg* is uniquely susceptible to lytic viruses in the *Haloferax* genus, potentially attributable to its lack of a CRISPR-Cas defense system [57].

HFTV1 induces cell lysis approximately 6 hours post-infection [58]. It possesses a dsDNA genome with 80 open reading frames (ORFs) and one viral tRNA (vtRNA) of unclear function [58]. Its entire virion has been resolved by cryo-EM [28], revealing tail fibers that likely mediate its recognition of the host S-layer [59]. Recently, Schwarzer et al. [58] characterized the transcriptional dynamics of HFTV1 and *Hg* during infection. They suggested potential post-transcriptional regulation via *cis*-acting antisense RNA (asRNA) binding complementarily to mRNA from the opposite strand to silence its translation [58, 60], although this remains to be validated experimentally. Additionally, they found that components of the *Hg* archaellum (archaeal equivalent to the flagellum) were down-regulated during infection, substantiated by decreased motility of infected cells. However, the translational dynamics of the HFTV1 infection cycle and the role of the vtRNA remain uncharacterized. Filling this gap would deepen our understanding of host-virus interactions in archaea at the molecular level. Furthermore, it would reveal the extent of post-transcriptional regulation in this system, laying the foundation to explore how this might be achieved in a host whose information processing machinery is predominantly eukaryotic.

Translational activity can be studied at high resolution and high throughput using ribosome profiling (Ribo-seq) [61]. Ribo-seq captures genome-wide ribosome density across all transcripts at a given time point, providing a snapshot of translation *in vivo*. When Ribo-seq and RNA-seq are applied to the same samples, this enables the calculation of translational efficiency (TE), a metric that can be used to decouple the effects of transcriptional and post-transcriptional regulation for a given gene [61, 62]. Ribo-seq is also more effective at detecting small or novel proteins than mass spectrometry [61], and, when combined with translation inhibitors, it can resolve features such as cryptic translation start sites [63–65]. Ribo-seq has been applied to several eukaryotic viruses [66–74], infrequently in phages [75, 76] and, to our knowledge, never before in archaeal viruses.

In this study, we characterized the dynamics of HFTV1 and *Hg* translation by applying Ribo-seq paired with RNA-seq throughout the infection cycle. We collected these data at high temporal resolution, capturing abrupt dynamics of gene expression particularly during early infection. We found evidence of post-transcriptional regulation for both *Hg* and HFTV1 genes; this included the late HFTV1 gene *gp17*, which was regulated in a manner that provided further support to the evolutionary trajectory of icosahedral viruses. The HFTV1 vtRNA co-sedimented with *Hg* ribosomal complexes, implicating it in translational regulation. Clustering analysis revealed evidence of co-regulation between host and viral genes, helping us detect a previously unannotated host fimbrial system that may mediate biofilm formation as an antiviral defense mechanism. Together, this work provides high-resolution insight into translation during viral infection at the evolutionary interface between bacteria and eukaryotes.

## RESULTS

### Transcription and translation persisted for both *Hg* and HFTV1 during infection

To investigate the dynamics of HFTV1 and *Hg* gene expression, we applied Ribo-seq paired with RNA-seq across the infection cycle (**Fig. 1A**), capturing both the abundance and the ribosome density of host and viral mRNA. Cell lysis occurred at 3.5-4 hours post-infection (p.i) at 42 °C (**Fig. S1**), so all samples were collected before 3.5 hours p.i (or 210 minutes p.i.). We collected material for Ribo-seq and RNA-seq at 0, 5, 15, 30, 60, and 180 minutes p.i (T0-T180) from the same cultures for 3 biological replicates (**Fig. 1B**). We also independently collected material at 150 and 200 min p.i (T150 and T200), the latter time point immediately preceding lysis. We obtained 9-28 million read pairs per Ribo-seq library (**Table S1**) and 8-33 million read pairs per RNA-seq library (**Table S2**). The native *Hg* plasmid pHGLR3 was absent in one biological replicate (**Fig. S2**), so we made appropriate adjustments to our analysis pipeline (see Methods). Viral features were reannotated with an automated pipeline as previously described [58, 77]. Furthermore, translation start sites were mapped by applying Ribo-seq to infected cells treated with harringtonine, a translation inhibitor that stalls ribosomes at initiation [78]. This enabled the revision of the 5’ boundaries of several viral ORFs (five extensions and three truncations) and the detection of a novel ORF encoding a hypothetical protein (**Fig. S3-S4** and **Table S3**). In total, our version of the viral genome contained 82 ORFs and one vtRNA. Reads were mapped to a merged host-virus genome for further analyses.

**Figure 1.**
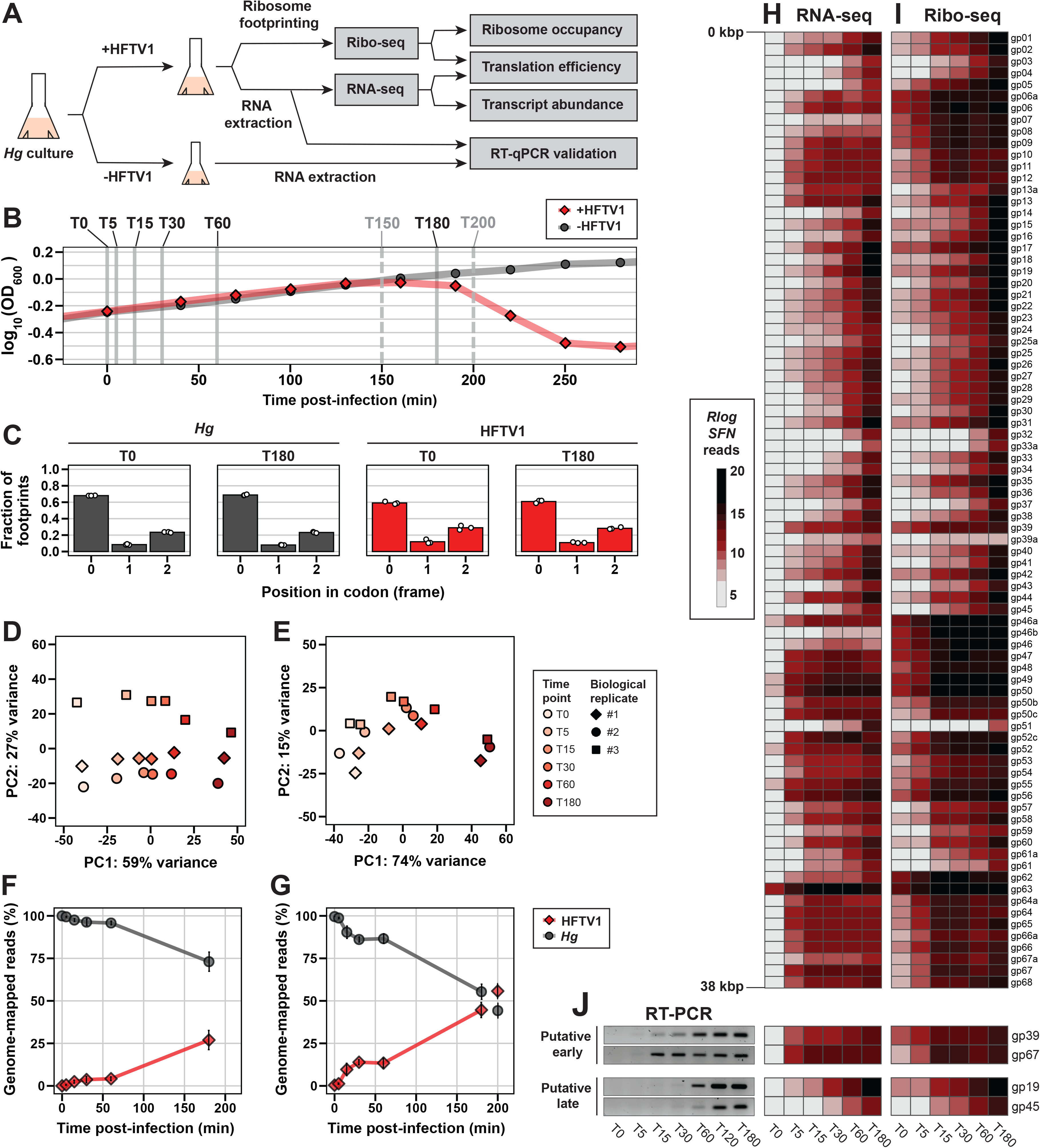
Global transcriptional and translational data across the HFTV1 infection cycle. **(A)** Schematic of experimental design. **(B)** Growth curve of *Hg* +/-HFTV1 annotated with post-infection sampling times. Cells grown to ORD600 at 42 °C, then infected with a multiplicity of infection (MOI) of 10. The paired RNA-seq/Ribo-seq time series (T0, T5, T15, T30, T60, and T180) is indicated by black text and solid vertical lines; additional sampling time points (T150 and T200) are indicated by grey text and dashed lines. RNA-seq/Ribo-Seq T0-T180 samples were collected for three biological replicates. Ribo-seq T150 and T200 samples were collected for two biological replicates. Data points (red diamonds and grey circles for +HFTV1 and - HFTV1, respectively) are growth curve data only and were collected independently of cultures used for RNA-seq/Ribo-seq. **(C)** Codon frame analysis showing 3 nt periodicity of the 5’ ends of elongating (27 nt) ribosome footprints at early (T0) and late (T180) time points. Reads mapped to *Hg* genes are in grey; reads mapped to HFTV1 genes are in red. Each point represents a biological replicate, and bar height is the mean of biological replicates; error bars were excluded since they were barely visible on some bars. **(D-E)** PCA of regularized log-transformed (*Rlog; DESeq2*), *sizeFactor*-normalized (*SFN; DESeq2*) reads of *Hg* and HFTV1 genes across time points and biological replicates for RNA-seq **(D)** and Ribo-seq **(E)**. **(F-G)** Percentage of all genome-mapped reads that mapped to *Hg* versus HFTV1 in RNA-seq **(F)** and Ribo-seq **(G)** datasets; T200 time points indicated as disconnected from T0-T180 data. **(H-I)** Heatmaps of *Rlog SFN* reads across all HFTV1 genes, ordered according to their position in the genome, for T0-T180 RNA-seq **(H)** and Ribo-seq **(I)**. **(J)** Expression of a subset of early and late HFTV1 genes across time points assessed by reverse transcription PCR (RT-PCR; left), RNA-seq (middle), and Ribo-seq (right). *Rlog:* regularized log-transformed (*DESeq2*). *SFN: sizeFactor-* normalized (*DESeq2*).

Ribosome footprints were examined at each time point for hallmarks of active translation.

The most abundant ribosome footprints were 24-27 nt long for both HFTV1- and *Hg-*mapped reads (**Fig. S5**), a length previously associated with elongating ribosomes in *Hv* [78, 79].

Ribosome density was highest at the first nucleotide of each codon for both HFTV1 and *Hg* ORFs (**Fig. 1C** and **Fig. S6**), and metagene plots of *Hg* ORFs showed three-nucleotide periodicity, with enriched ribosome density in coding regions versus noncoding regions (**Fig. S7**). This periodicity has previously been used in *Hv* to distinguish genuine ribosome density from artefactual MNase degradation [83]. These metrics confirmed that our data captured the activity of elongating ribosomes. HFTV1 metagene plots were noisier and less conclusive (**Fig. S7**), which we attributed to the smaller number of HFTV1 ORFs (<100) relative to *Hg* (∼4,000) rather than to read quality. Furthermore, polysome profiles collected independently from Ribo-seq data showed abundant polysomes at 30, 60, 120, and 180 min p.i. (**Fig. S8**). Together, these results indicated an active translation system at all time points, suggesting that mRNA from both HFTV1 and *Hg* was translated throughout the infection cycle.

Principal component analysis of combined HFTV1 and *Hg* expression data in all samples showed that PC1 accounted for most variance in RNA-seq (**Fig. 1D**, 59%) and Ribo-seq (**Fig. 1E**, 74%), stratifying samples in temporal order from T0 to T180 and clustering replicate samples. Over time, genome-mapped reads increased for HFTV1 and decreased for *Hg* in both RNA-seq (**Fig. 1F**) and Ribo-seq (**Fig. 1G**); by T180, the proportions of *Hg-* and HFTV1-mapped reads were ∼75% and ∼25% in RNA-seq and roughly 50% and 50% in Ribo-seq.

To assess the agreement between RNA-seq and Ribo-seq at the level of single genes, we looked at normalized read counts from both datasets along the HFTV1 genome. Consistent with the temporal organization of viral gene expression (e.g., early vs. late), we found that large “blocks” of genomically adjacent HFTV1 genes followed consistent temporal expression patterns in RNA-seq (**Fig. 1H**) and Ribo-seq (**Fig. 1I**). Some blocks of viral genes were expressed very early during infection (e.g., *gp05-gp13*, *gp46a-gp50c*, *gp52c-gp56*, and *gp62-gp68*), whereas others were expressed later (e.g., *gp14-gp30*, *gp57-p61*). We validated a subset of these genes qualitatively by RT-PCR (**Fig. 1J**). Notably, a handful of viral genes had ribosome density that was disproportionately higher than transcript abundance (e.g., *gp07, gp46b,* and *gp62*), but these comprised a minority of the genome. Overall, our data captured consistent temporal expression patterns for the HFTV1 and *Hg* genomes during infection.

To distinguish between the effects of transcriptional and post-transcriptional regulation, we calculated translation efficiency (TE) by dividing normalized Ribo-seq reads by normalized RNA-seq reads for each gene. For all time points, most *Hg* genes (96.6-98.2%) exhibited translation efficiency between 0.1 and 10 (**Fig. S9**). Only 30-55 genes per time point fell outside this range, many encoding hypothetical or uncharacterized proteins, with no functional trends among genes with meaningful predicted functions. We therefore restricted further investigations of TE to genes with |TE| > 10 that emerged as genes of interest through other analytical approaches.

Despite the broad similarities between our RNA-seq and Ribo-seq datasets, we focused on Ribo-seq for subsequent analyses to capture any nuanced translation-level changes and used RNA-seq to verify whether trends were shared across both datasets or unique to one.

Together, these approaches centered our analysis on the translational landscape of infection.

### Clustering analysis revealed shared expression patterns among genes with differential ribosome density

To identify genes and processes that play a critical role during the infection cycle, we used a likelihood ratio test (LRT) with *DESeq2* [80] to detect genes with differential ribosome density (dRibo). This method tested whether variation in time accounted for variation in normalized read counts per gene, in contrast to the pairwise Wald test that compares only two conditions at a time. We selected a stringent significance threshold to provide high confidence in the association between expression pattern and time (BH-adjusted p-value < 10e-5). With this approach, we identified 1,721 dRibo genes (**Fig. 2A**, **Table S4)** independent of their magnitude of change between any two time points. This accounted for ∼44% of all tested *Hg* genes and 100% of all tested HFTV1 genes. Similar statistics were obtained for genes with differential transcript abundance (dRNA; **Table S5**). This showed that almost half of the *Hg* genome was responsive to the time course of infection and that all viral genes were expressed.

**Figure 2.**
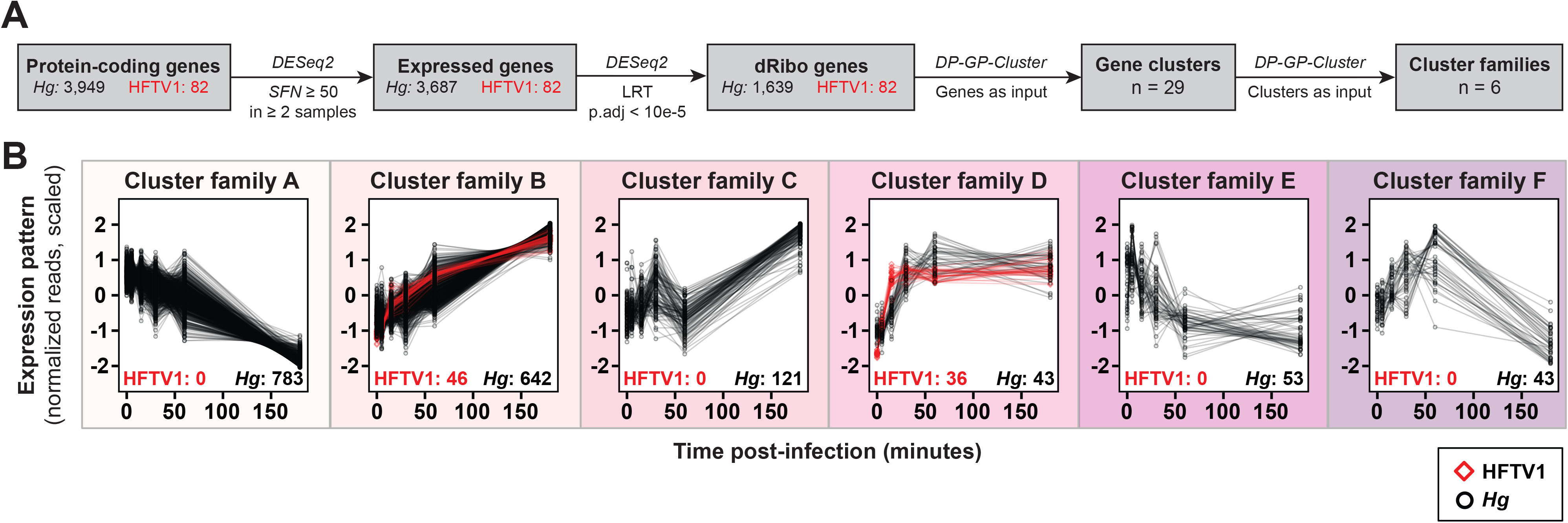
Clustering revealed shared expression patterns among genes with differential ribosome density (dRibo). **(A)** Analysis pipeline to identify and cluster dRibo genes. Only protein-coding genes were used for differential expression analysis. **(B)** Cluster families produced by clustering mean expression trajectories of gene clusters; visualized using scaled *Rlog SFN* reads. The number of genes within each cluster family are indicated in black for Hg and red for HFTV1. CDS: Coding sequence. *Rlog:* regularized log-transformed (*DESeq2*). *SFN: sizeFactor-*normalized (*DESeq2*).

To detect common trajectories of differentially expressed genes, we clustered all dRibo genes by temporal expression pattern using *DP_GP*_*Cluster* [81], an unsupervised software that minimizes user biases and performs well with temporal expression data. Expression patterns were mean-centered and scaled so that the overall expression pattern would be considered without interference from the magnitude of expression change. This generated 29 clusters of dRibo genes (**Fig. S10-S11**). We consolidated these clusters by applying *DP_GP_Cluster* again to the mean trajectory of each cluster rather than to the trajectory of each gene. This meta-clustering step did not produce *bona fide* gene clusters, so we instead referred to the output as “cluster families”. Both clustering steps were performed for dRNA genes and produced qualitatively similar clusters and cluster families (**Fig. S12-S14**).

Our two-step clustering process generated six cluster families of dRibo genes (**Fig. 2B** and **Table S6**). These cluster families varied dramatically in size, ranging from 43 to 783 genes. HFTV1 genes were present in only Cluster Families B (46 HFTV1 genes) and D (36 HFTV1 genes). Most dRibo genes were either gradually down-regulated (Cluster Family A) or gradually up-regulated (Cluster Family B). The remaining cluster families followed more dramatic trends and accounted for fewer genes (Cluster Families C-F). We also calculated log2 fold-change (L2FC) values at each time point relative to T0 for all genes (**Fig. S10-S11** and **Table S6**).

Having grouped all genes in a temporally meaningful way, we then analyzed the functions of the constituent genes in each cluster family - first for HFTV1 genes, then for *Hg* genes - to understand the stratification, interplay, and regulation of viral and host processes over time.

### HFTV1 genes follow early and late temporal expression patterns

All HFTV1 genes were up-regulated (**Fig. 2B-C**, Cluster Families B and D). Most of them (95%) could not be automatically assigned to arCOG-based functional categories [82], so we assigned them manually to custom functional categories (**Fig. S15**). HFTV1 genes in Cluster Family D were sharply up-regulated at early time points (T15 vs. T0 mean L2FC = 5.3 ± 0.51 SD, n = 36), then they plateaued and remained stably expressed through T180 (T180 vs. T0 mean L2FC = 6.1 ± 1.0 SD, n = 36). These genes comprised putative transcription factors and core DNA replication and recombination machinery (**Fig. S15**), including a PCNA clamp, a DNA helicase, and a RecT-like ssDNA annealing protein. In phages, transcription factors are most abundant in early and middle gene classes, and DNA replication machinery is most abundant in middle gene classes [83]. Since our clustering approach produced only two viral gene classes, we suggest that the viral genes in Cluster Family D comprise the early genes of HFTV1. In contrast to these early genes, the HFTV1 genes in Cluster Family B increased gradually in expression (**Fig. 2B**). While they were initially up-regulated to a lesser degree than early genes (T15 vs. T0 mean L2FC = 2.9 ± 1.4 SD, n = 46), they vastly exceeded early genes by late stages (T180 vs. T0 mean L2FC = 9.5 ± 1.0 SD, n = 46). These genes were associated with characteristically late viral processes: virion structure, packing, and assembly (**Fig. S15**) [83]. The major capsid protein and the capsid stabilization protein showed strong induction between T60 (T60 vs. T0 L2FC = 4.5-4.9) and T180 (T180 vs. T0 L2FC = 10.5-11.1). This gene set also included several transcription factors, the ribosome-dependent endonuclease RelE, and a phosphoadenosine phosphosulfate reductase, a metabolic enzyme involved in sulfur assimilation [84]. Based on their functions and expression patterns, we suggest that HFTV1 genes in Cluster Family B comprise the late genes of HFTV1.

### The HFTV1 scaffold protein was translated from an in-frame internal ORF of the prohead protease gene *gp17*

During our analysis of HFTV1 genes, we identified a discrepancy between RNA-seq and Ribo-seq coverage along the late gene *gp17* (**Fig. 3A**). While transcript abundance showed no distinct coverage bias, ribosome density seemed highly enriched in the 3’ region of the transcript compared to the 5’ region. Based on the region of high ribosome density, we divided *gp17* into a 5’ region (nucleotide positions 1-876) and a 3’ region (nucleotide positions 877-1,443) and assessed the cumulative reads per million in each region at T180. Transcript abundance was significantly lower in the 3’ region compared to the 5’ region by a factor of 2.3 (**Fig. 3B**; p-value < 0.05). In contrast, ribosome density was significantly higher in the 3’ region compared to the 5’ region by a factor of 3.1 (**Fig. 3C**; p-value < 0.005). By calculating translation efficiency for each region, we found that the 3’ region had higher translation efficiency than the 5’ region throughout infection (**Fig. S16A**). Together, these data suggested that *gp17* was post-transcriptionally regulated.

**Figure 3.**
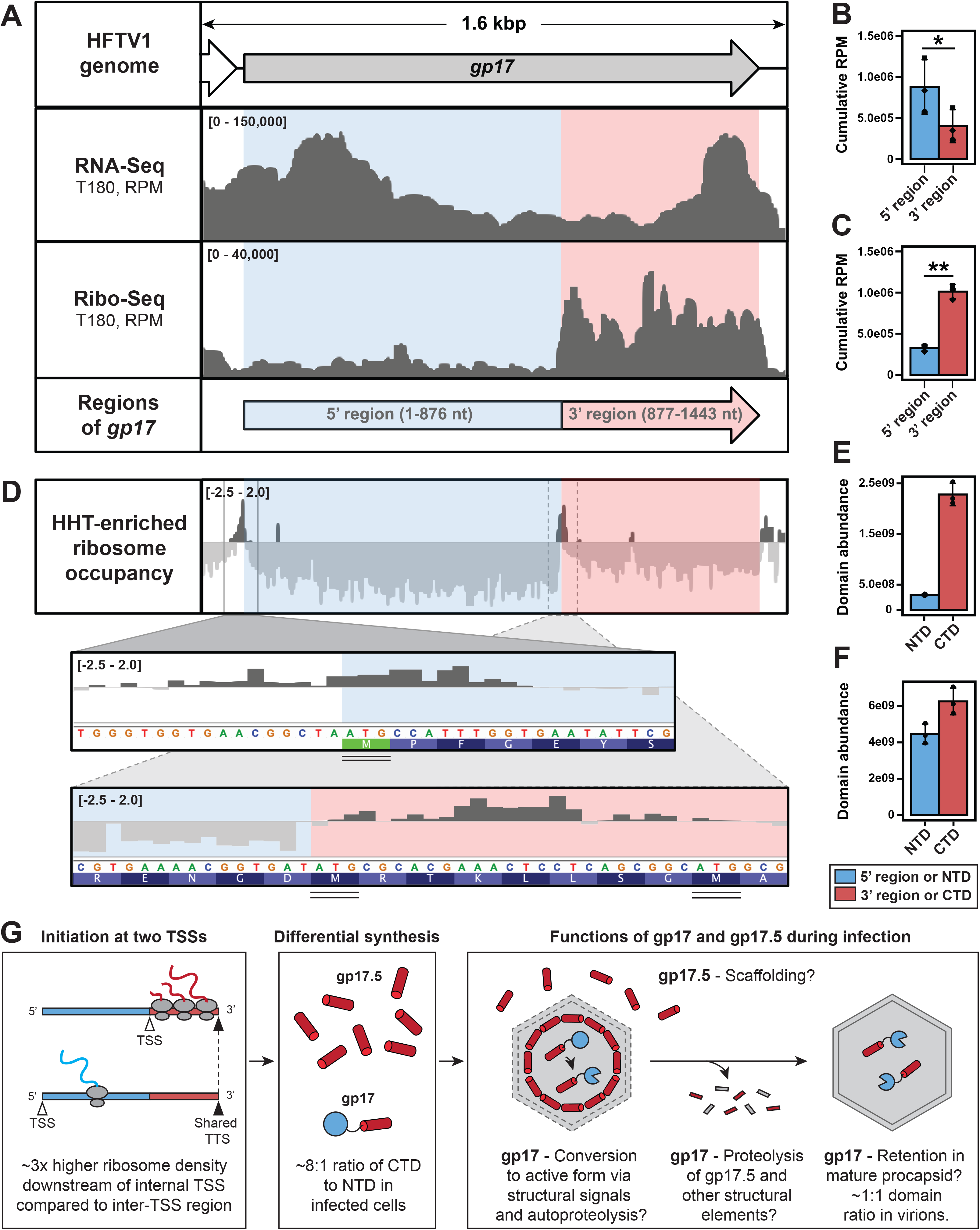
The HFTV1 scaffold protein was translated from an in-frame internal ORF of the prohead protease gene *gp17*. **(A)** HFTV1 gene *gp17* (from Fig. 2, Cluster Family B) and corresponding RNA-seq and Ribo-seq coverage at T180, expressed as RPM. Boundaries of 5’ and 3’ regions defined by Ribo-seq coverage are indicated at the bottom of the panel; 5’ region is in red, 3’ region is in blue. **(B)** Cumulative RNA-seq RPM by *gp17* region (unpaired two-tailed Student’s T-test, three biological replicates, *p-value < 0.05). **(C)** Cumulative Ribo-seq RPM by *gp17* region (unpaired two-tailed Student’s T-test, 3 biological replicates, **p-value < 0.005). **(D)** Ribo-seq data from HHT-treated cultures normalized to paired data from untreated-culture and transformed at each nucleotide position as log2([+HHT RPM + 1]/[-HHT RPM + 1]). Magnified regions focus on the two highest peaks of HHT-induced ribosome density along *gp17*: the canonical TSS and an internal site. The corresponding nucleotide sequence and amino acid sequence of the *gp17* open reading frame are shown for each magnification, with AUG start codons double-underlined. **(E-F)** Abundance by LC-MS/MS of the gp17 N-terminal domain (NTD), derived from the 5’ region, and the gp17 C-terminal domain (CTD) derived from the 3’ region, in **(E)** HFTV1-infected *Hg* at T180 and **(F)** Expression of the gp17 domains analyzed by LC-MS/MS in HFTV1-infected *Hg* at T180. Left: purified virions. **(G)** Model of the regulation, approximate ratios, and function of the prohead protease and scaffold proteins in HFTV1. TSS: Translation start site. TTS: Translation termination site. All genome tracks in **A** and **D** represent one of three biological replicates. All error bars represent SD. T180: 180 minutes post-infection. RPM: Reads per million. HHT: Homoharringtonine. *SFN: sizeFactor-*normalized reads (via *DESeq2*). NTD: N-terminal domain of gp17. CTD: C-terminal domain of gp17.

The *gp17* ORF contained several in-frame AUGs, any one of which could serve as an internal translation start site (TSS) for an N-terminally truncated form of the protein. To test this, we performed Ribo-seq on HFTV1-infected cells treated with harringtonine (HHT), which arrests ribosomes at initiation and causes ribosomes to accumulate at TSSs in archaea and eukaryotes (**Fig. S3-S4**) [63, 78, 85]. Normalized HHT-treated data revealed two initiation-associated peaks in *gp17*: one at the canonical 5’ TSS, and another in the 3’ half of *gp17* overlapping nucleotide positions 877-899 (**Fig. 3D**), directly preceding the region with high ribosome density seen in standard Ribo-seq (**Fig. 3A**). Both peaks were proximal to in-frame AUGs with no intervening stop codons (**Fig. 3D**, magnifications), indicating that *gp17* contained two TSSs capable of producing both the full protein and an N-terminally truncated form from the same reading frame. Previously, asRNA was proposed to regulate gene expression in HFTV1 post-transcriptionally based solely on abnormal RNA-seq coverage patterns [24], so we examined *gp17* asRNA coverage. We found no evidence to support the role of asRNA regulation along *gp17* (**Fig. S16B-C**). The mechanism of *gp17* TSS selection remains unknown.

We then used mass spectrometry to validate the differential ribosome density of the 5’ and 3’ regions of *gp17* (**Fig. S17** and **Table S7**). For our analysis, we split gp17 into an N-terminal domain (NTD, residues 1-292, corresponding to the 5’ region) and a C-terminal domain (CTD, residues 293-480, corresponding to the 3’ region). Because the comparison of domain abundance was semi-quantitative, we did not test for significance; nonetheless, at T180, the CTD was 7.8x more abundant than the NTD (**Fig. 3E**), indicating that some of the CTD was synthesized independently of the NTD and validating our Ribo-seq data (**Fig. 3C**). However, it remained unclear how increased synthesis of the CTD influenced gp17 function.

The protein encoded by *gp17* is annotated as a “P2 gpO-like scaffolding/protease protein” (from RefSeq assembly GCF_004208775.1; submitted Jan 22, 2019), presumed to be a fused form of the capsid scaffold and the prohead protease [16]. In bacteriophage P2, gpO features a CTD that scaffolds the capsid during assembly and a proteolytic NTD that matures the capsid and other structural components [86]. Although gpO is synthesized as a single full-length protein, its NTD later autoproteolytically cleaves itself from the CTD; the NTD remains inside the mature capsid while the CTD is degraded. We found minimal sequence or structural homology between gp17 and gpO (19% sequence ID, 19% structure ID), but we speculated that their function was still conserved. We hypothesized functional conservation between gpO and gp17 would result in high NTD abundance and low CTD abundance of gp17 in mature capsids, which we tested by analyzing the protein content of purified virions by mass spectrometry (**Fig. S17-S18** and **Table S7-S8**). Contrary to our hypothesis, both the NTD and CTD were abundant in purified virions (**Fig. 3F**), with the CTD only 1.4x more abundant than the NTD, much closer to a 1:1 ratio of NTD and CTD than in infected cells. These results suggested that the truncated form of gp17 (CTD only) was more abundant in infected cells, whereas the full form of gp17 (NTD and CTD in roughly equal ratio) was more abundant in the mature capsid. Thus, contrary to its annotation, gp17 is functionally unlike the phage P2 protein gpO. Instead, it seems more similar in function and gene organization to "nested" protease-scaffold proteins found in certain dsDNA bacterial and eukaryotic viruses [87].

### The HFTV1 vtRNA was expressed during infection and associated with host ribosomes

The HFTV1 genome encodes one vtRNA, which is predicted to decode the threonine codon ACG. This vtRNA^Th^_rCGU_ contains a putative 5-nt intron within the anticodon region, a feature consistent with archaeal tRNAs [88]. To assess whether the predicted transcript could adopt a tRNA-like conformation, we modelled its secondary structure using RNAFold [89]. The predicted secondary structure formed a tRNA-like fold, although it deviated from the canonical tRNA secondary structure due to the presence of the intron (**Fig. 4A**, left). Removal of the putative intron generated a mature anticodon loop (**Fig. 4A**, right). To assess whether the vtRNA was expressed during infection, we performed Northern blot analysis of total RNA from *Hg +/-* HFTV1 using a probe specific to the 5’ region of the vtRNA and to *Hg* tRNA^Thr^_CGU_, which was used as a control. We detected a vtRNA-specific signal as early as 60 min p.i. (**Fig. 4B**) and an accumulation of two distinct RNA species from 120 min p.i. onward.

**Figure 4.**
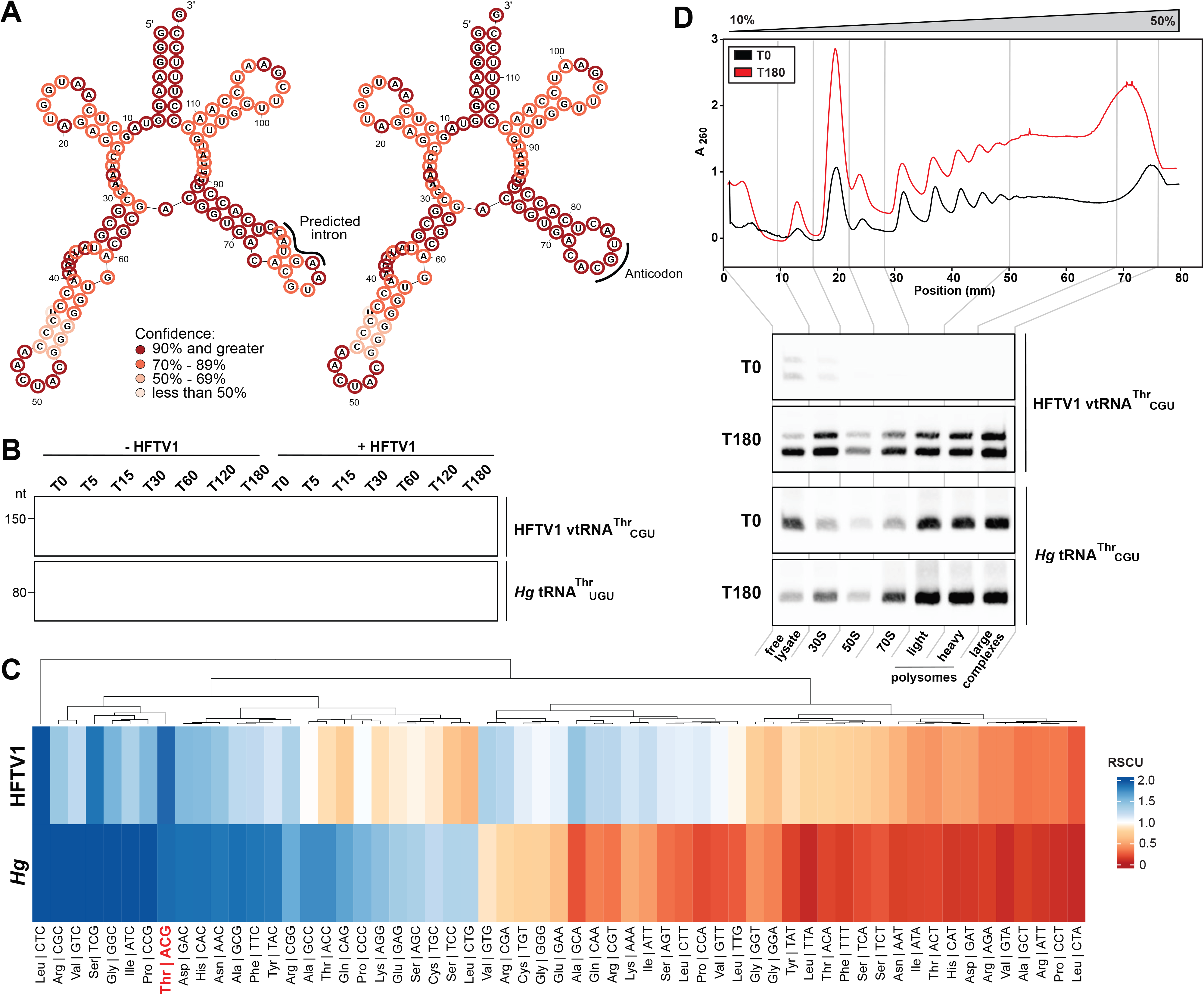
The HFTV1 vtRNA was expressed during infection and co-sediments with translating ribosomes. **(A)** 2D structure prediction of vtRNA^Thr^_CGU_ with intron (left) and without intron (right). Predicted intron (5 nt) and anticodon are marked with line. **(B)** Northern blot analysis of HFTV1 vtRNA^Thr^_CGU_ (top) and *Hg* vtRNA^Thr^_CGU_ (bottom) expression +/-HFTV1 infection. **(C)** Relative synonymous codon usage (RSCU) in the genomes of HFTV1 (top) and *Hg* (bottom). The Thr codon ACG decoded by vtRNA^Thr^_CGU_ is highlighted in red. **(D)** Polysome profiles along a 10-50% sucrose gradient (top) and Northern blot analysis of corresponding gradient fractions (bottom) showing co-sedimentation of HFTV1 vtRNA^Thr^_CGU_ and *Hg* tRNA^Thr^_CGU_ with fraction contents at 0 and 180 minutes post-infection.

Viral tRNAs have been proposed to compensate for differences in synonymous codon usage between the viral and host genomes [90, 91]. To assess whether the HFTV1 vtRNA could serve such a role, we compared relative synonymous codon usage (RSCU) values across all annotated CDSs of HFTV1 and *Hg*. We found no evident enrichment of the ACG threonine codon in the virus genome relative to the host (**Fig. 4C**).

We then investigated whether the putative vtRNA associates with the *Hg* translational machinery during HFTV1 infection. For HFTV1-infected *Hg* at 0 and 180 min p.i., we isolated monosomes and polysomes following ultracentrifugation of host ribosomes on a sucrose gradient and extracted RNA from individual fractions. We probed for tRNAs in individual or pooled fractions by Northern blotting with the probe specific to the 5’ vtRNA region, with *Hg* tRNA^Thr^_CGU_ as a control for host tRNA distribution across the polysome profile. Although a vtRNA-specific signal was detected in all analyzed fractions, the vtRNA was clearly enriched in fractions corresponding to the 30S subunit and actively translating polysomes (**Fig. 4D**). This distribution resembled that of *Hg* tRNA^Thr^_CGU_, indicating that the HFTV1 vtRNA is associated with host ribosomal complexes during infection.

### Infection with HFTV1 reshaped the translational landscape of *Hg*

After examining viral gene expression, we focused on the consequences of infection for *Hg*. Host dRibo genes varied across a range of L2FC values, with a maximal down-regulation of L2FC = -5.7 (T180 vs. T0) and a maximal up-regulation of L2FC = 5.8 (T180 vs. T0) (**Fig. S13**). Genes from all functional categories were dRibo (**Fig. 5A**). Functional categories with the largest dRibo proportions corresponded to translation and ribosome structure/biogenesis (arCOG J, 76% dRibo, n = 125) and to nucleotide transport and metabolism (arCOG F, 77% dRibo, n = 58). The smallest dRibo proportion pertained to cell motility (arCOG N, 18% dRibo, n = 8). Proportions of differentially expressed genes within each functional category were roughly consistent across Ribo-seq and RNA-seq (**Fig. S19**).

**Figure 5.**
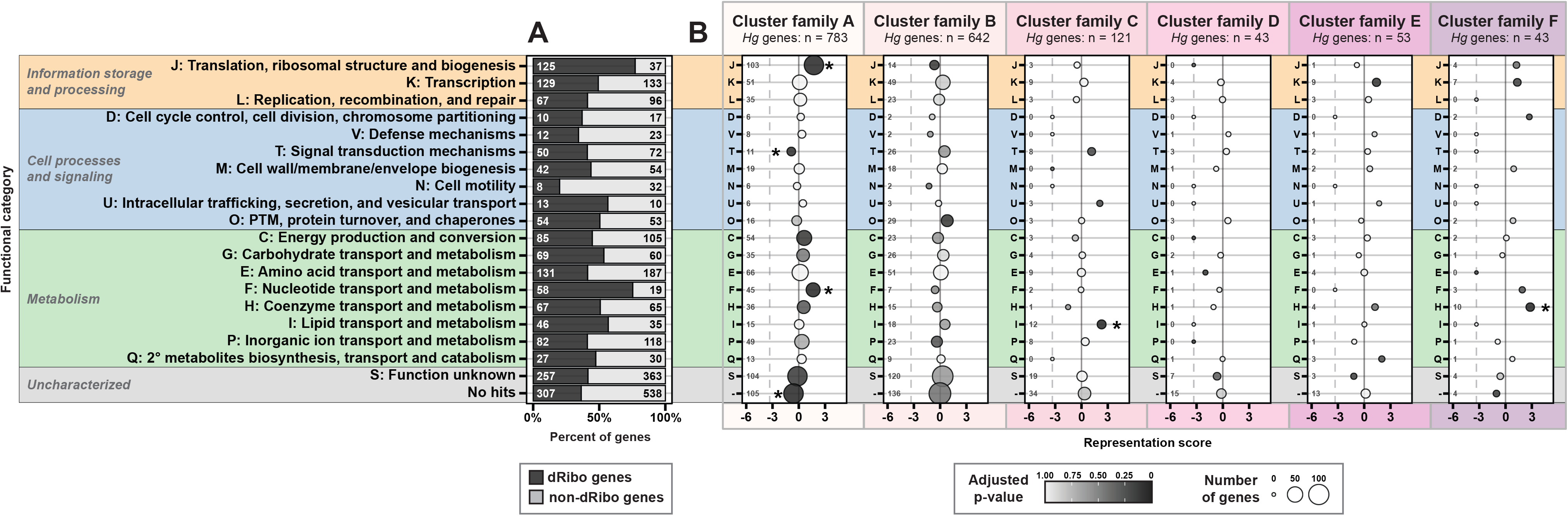
Functions of *Hg* genes with differential ribosome density per cluster family. **(A)** Proportions of dRibo and non-dRibo *Hg* genes per arCOG functional category. **(B)** Representation analysis of the functional categories of *Hg* genes in each cluster family. Representation is relative to the number of genes per functional category in the *Hg* genome. Representation scores were calculated in the following manner: score = log2[([no. of genes for given category in cluster family]/[expected no. of genes for given category in cluster family based on reference genome] + 0.1)]. Representation score = 0 (solid vertical grey line) indicates that a functional category was represented in a cluster family to the same degree as in the genome, score > 0 indicates the category was over-represented, and score < 0 indicates the category was under-represented. Some categories are significantly over- or under-represented, indicated by (*), with BH-adjusted p-value < 0.05. Due to the score calculation, points along the dashed vertical grey line correspond to functions that had no representative genes in a given cluster family. The arCOG category "No hits" indicates genes in the *Hg* genome that could not be assigned to a functional category. dRibo: differential ribosome density. For cluster family expression patterns, see Fig. 2B.

#### Core information processing machinery is gradually down-regulated during infection

Almost half of all dRibo host genes (783/1,639 = 48%) were down-regulated during infection (**Fig. 5B**, Cluster Family A; for corresponding expression patterns, see **Fig. 2B**). This accounted for 94% of translation-related (arCOG J) genes in the *Hg* genome, including all ribosomal proteins, translation initiation factors aIF6 and aIF2, elongation factors, all tRNA ligases, and several tRNA modification and maturation enzymes (T180 vs. T0 mean L2FC = -1.52 ± 0.64 SD, n = 103). Other down-regulated components of information processing machinery included the DNA-directed DNA polymerase, all RNA polymerase subunits, and transcription elongation factors (**Fig. 5B**, Cluster Family A, arCOG K and L). Nucleotide metabolism was similarly down-regulated (**Fig. 5B**, Cluster Family A, arCOG F), along with the production of cobalamin, a coenzyme associated with nucleotide biosynthesis (Cluster Family F, arCOG H). This indicated that nucleotide production and most core machinery of the information processing system was slowly down-regulated during infection.

#### Universally conserved translation initiation factors were stably expressed

Since most translation machinery components were down-regulated, we explored the exceptions to this trend. Select up-regulated genes related to translation included three translation initiation factors: aIF1, aIF5B, and aIF1A (**Fig. S20**). Changes in the expression levels of these factors were modest (T180 vs. T0 L2FC values = 0.55-0.93), showing that their expression increased or was at least sustained during late infection. In contrast, aIF2 subunits and aIF6 were clearly down-regulated (T180 vs. T0 L2FC values: aIF2, -1.76 ± 1.0 SD; aIF6, -1.40). Notably, the functions of aIF1, aIF5B and aIF1A are universally conserved across archaea, bacteria, and eukaryotes [11]. In contrast, the down-regulated factors aIF2 and aIF6 are functionally conserved only between archaea and eukaryotes [11, 92]. This suggests a link between the evolutionary conservation of translation initiation factors and their expression patterns during infection.

#### A widespread transcriptional network links basal and specific transcription factors across temporal gene groups

Host transcription factors (TFs) were found across all cluster families, showing diverse and widespread expression changes during infection (**Fig. 5B**, arCOG K; for corresponding expression patterns, see **Fig. 2B**). Differentially expressed TFs included basal TFs (TATA-binding proteins and several copies of TFB) and over 70 specific regulators from families such as ArsR/SmtB, TrmB, Lrp/AsnC, IclR, PadR, XacR, and MarR, suggesting infection engaged a broad transcriptional network. TFs in Cluster Family E showed the most dramatic response of any *Hg* genes, spiking at T5 (mean L2FC of 3.5 ± 0.38; T5 vs. T0, n = 4) before returning to baseline levels by T15. Three of these TFs (HfgLR_RS02315, RS02290, and RS17090) also showed unusually high translation efficiency (TE > 10) at T5, though the TE of only one (HfgLR_RS17090) was sustained throughout infection, suggesting that the immediate response of specific TFs might be shaped by post-transcriptional regulation (**Fig. S8**). Overall, TFs or putative TFs made up 20% (n=11) of Cluster Family E, together with stress response genes (universal stress proteins and a methyltransferase of a putative R-M system). These findings suggest that Cluster Family E reflects a rapid, transient induction of defense and stress response pathways that is then quickly aborted.

#### Protein quality control machinery remained active during late infection

Genes associated with protein folding, post-translational modification, and protein turnover (arCOG O) were not significantly over-represented in any cluster family but were nonetheless predominantly up-regulated at late time points (**Fig. 5B**, Cluster Family B; for corresponding expression pattern, see **Fig. 2B**). Chaperones, specifically subunits of the thermosome, accounted for the largest changes in expression level (T180 vs. T0 L2FC = 2.1-3.5, n = 3). Other genes included proteasome-activating nucleotidases (T180 vs. T0 L2FC = 1.3-1.4, n = 2), proteasome subunits (T180 vs. T0 L2FC = 0.52-0.56, n = 2), and several thioredoxins or other thioredoxin-like proteins predicted to reduce sulfide bonds as a form of PTM (T180 vs. T0 L2FC = 0.44-1.52, n = 9). All of these genes were co-regulated with late viral genes, following a gradually increasing trajectory over time. This could indicate increased protein misfolding or proteolytic turnover at late infection stages and an attempt by the cell to regain homeostasis.

#### Host amino acid import increased prior to lysis

To capture expression changes in the hour before cell lysis, we used additional Ribo-seq data from T150 and T200, analyzed in a standard pairwise fashion (*DESeq2* Wald test, |L2FC| > 1.5, BH-adjusted P-value < 0.05). Few HFTV1 genes changed between these time points, but *Hg* genes showed substantial shifts: 75 genes up-regulated and 100 genes down-regulated (**Fig. S21**). Notably, almost 40% (n = 23) of up-regulated *Hg* genes encoded complete ABC-type oligopeptide import systems (**Fig. S21**), including nucleotide-binding proteins, transmembrane proteins, and substrate-binding proteins (mean L2FC = 2.03 ± 0.45 SD) [93]. This suggested that *Hg* attempted to import substantial amounts of amino acids from the extracellular environment just before lysis, potentially in response to amino acid starvation.

### A Tat-dependent host fimbrial system is co-regulated with early viral genes

In the same Cluster Family as early viral genes (Cluster Family D, **Fig. 6A**) we identified three genes (HfgLR_RS13645, 13650, and 13655, hereafter described as 13645-13655) that exhibited some of the highest L2FC values out of all early *Hg* genes (T30 vs. T0 L2FC = 3.0-3.3). The proteins encoded by 13645-13655 had no substantial sequence-based homology to characterized proteins, but structural prediction revealed that they were similar to subunits of Type IV pilin-like fimbriae (**Fig. 6B**). All three proteins encoded signal peptides at their N-termini; furthermore, the signal peptides of the 13645 and 13655 proteins contained twin-arginine translocation (Tat) export motifs (**Fig. 6B** and **Fig. S22**) [94].

**Figure 6.**
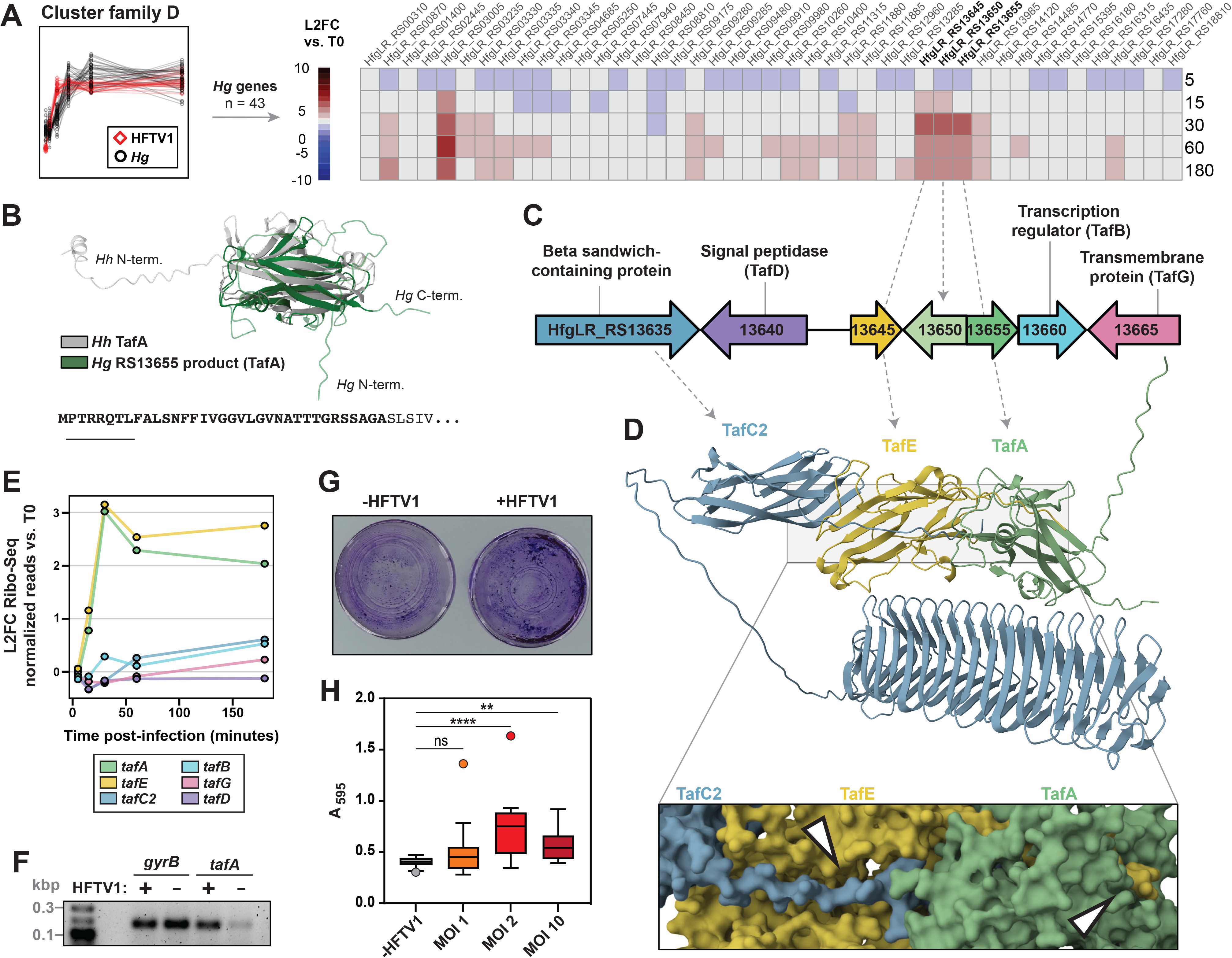
A Tat-dependent fimbrial system (Tafi) of *Hg* was co-regulated with early HFTV1 genes. **(A)** Cluster Family D (depicted as in Fig. 2B) and heatmap of expression patterns for all *Hg* genes from Cluster Family D, depicted as L2FC *SFN* reads relative to T0. **(B)** Structural alignment of *Hg* HfgLR_RS13655 product (TafA) and *Hh* Tafi subunit TafA (11% ID); structures predicted by AF2 and aligned using the PDB web tool [113]. The amino acid sequence shows the first 40 N-terminal residues of the HfgLR_RS13655 product, with signal peptide in bold and Tat signal underlined. **(C)** Genomic context of HfgLR_RS13645-13655, with labeled homologs of Tafi structural and auxiliary components. **(D)** AF2-predicted multimer of the HfgLR_RS13635, 13645, and 13655 products; magnification shows predicted donor strand exchange, with arrows pointing at donor strand from one subunit stabilizing another subunit. **(E)** Expression patterns of *Hg* Tafi components, colored as in **(C)**; depicted as L2FC *SFN* reads relative to T0. **(F)** RT-PCR of reference gene *gyrB* and *tafA* (HfgLR_RS13655) in HFTV1-infected versus mock-infected *Hg* at 30 minutes post-infection (plain media used for mock-infection); representative of three biological replicates (see **Fig. S27** for RT-qPCR data). **(G)** Photographs of *Hg* +/-HFTV1 stained with crystal violet, 180 minutes post-infection. Cells were infected in liquid medium with multiplicity of infection (MOI) = 10, then grown statically at 42 °C. **(H)** Quantitation of crystal violet-based biofilm assay of *Hg* +/-HFTV1; +HFTV1 conditions labeled based on MOI used for infection. T0: 0 minutes post-infection. Tafi: Tat-dependent fimbriae. RT-PCR: reverse transcription PCR. RT-qPCR: quantitative RT-PCR. L2FC: Log2 fold-change. *SFN: sizeFactor-* normalized reads (via *DESeq2*).

Within the ∼6 kbp region surrounding the 13645-13655 loci, we found homologs to several components of an archaeal-like Tat-dependent fimbrial (Tafi) system, including a transmembrane transcription regulator (TafB), a signal peptidase (TafD), and a transmembrane protein of unknown function (TafG) [95, 96] (**Fig. 6C**). There was initially no apparent gene nearby that could encode a tip with adhesin-like properties, but locus 13635 encoded a structural homolog to the *Natrinema* putative Tafi anchor TafF2 [96], sharing an N-terminal alpha-helix, a beta sandwich domain, and a C-terminal beta-helix domain (10 coils in *Natrinema*, 16 coils in *Hg*) (**Fig. S23B,D**). However, while the N-terminal alpha-helix was previously reported for *Natrinema* TafF2 as a transmembrane domain [96], we predicted with SignalP [97] that this helix was a Sec signal peptide for both *Natrinema* TafF2 and the product of *Hg* locus 13635 (**Fig. S23A,C**). Furthermore, the extended beta-helix resembled proteins with adhesin domains from bacteria [98, 99] and phages [100] (**Fig. S23F-G**). We therefore suspect that both *Natrinema* TafF2 and the protein of *Hg* locus 13635 function as Tafi tips rather than anchors.

Overall, except for the membrane anchor, we identified putative homologs for all structural and auxiliary Tafi components near the 13645-13655 loci.

All pairwise combinations of the 13635, 13645, and 13655 gene products were predicted to form high-confidence heterodimers (**Fig. S24A**; pLDDT > 80, pTM > 0.55, ipTM > 0.80), with 13645 and 13655 also forming high-confidence homodimers (**Fig. S24B**; pLDDT > 85, pTM > 0.85, ipTM > 0.85). The 13650 product dimerized with itself but did not form multimers with other components, so we excluded it from further analyses. At the heterotrimeric level, the remaining three structural proteins assembled in the order of 13635-13645-13655, with 13635 at the tip preferentially interacting with 13645 (**Fig. 6D** and **Fig. S24C**). Donor strand exchange was evident at both interfaces (**Fig. 6D**, inset), similar to Tafi in other haloarchaea [95, 96].

These data suggest the following identities: 13635 encodes a novel Tafi tip protein, 13645 encodes the adapter pilin TafE, and 13655 encodes the major pilin TafA. We propose to name the novel Tafi tip protein TafC2, given its distinct features relative to other systems [95, 96]. The pilin subunits TafA (13655) and TafE (13645) were highly upregulated at T30 (**Fig. 6E**). In contrast, the tip protein TafC2 was modestly up-regulated (L2FC = 0.60, T180 vs. T0), while the other auxiliary components (TafD, TafB, and TafG) were not dRibo during infection. The induction of *tafA* by viral infection was experimentally validated at T30 by reverse-transcription quantitative PCR (RT-qPCR; **Fig. 6F** and **Fig. S25**).

Consistent with its Tafi system, *Hg* encodes three components of the requisite Tat export pathway [101, 102]: TatD (HfgLR_RS11395), TatA (HfgLR_RS05905), and TatC (HfgLR_RS00905). All three components were dRibo during infection (**Table S6**), and *tatA,* which encodes the protein for Tat pore formation, consistently had some of the highest TE values of all *Hg* genes (**Fig. S9**, TE = 25.70 ± 9.28 SD, T0 to T180).

In other haloarchaea, Tafi play a role in biofilm formation [96]. Considering the co-regulation of the *Hg* Tafi system with early HFTV1 genes, we hypothesized that viral infection would induce or enhance biofilm formation in *Hg.* We tested this under infected and uninfected conditions with an assay that used crystal violet to stain biofilm. Indeed, we saw qualitatively that infection caused cells to retain more crystal violet when using a multiplicity of infection (MOI) of 10 (i.e., infecting with 10 virions per cell) (**Fig. 6G**). Quantification showed that A595 was significantly higher during infection with both MOI 2 and MOI 10 compared to uninfected controls (**Fig. 6H**).

## DISCUSSION

Archaeal host-virus interactions are poorly understood at the molecular level. Here we advance our understanding of this subject by characterizing the translational dynamics of HFTV1 and its archaeal host *Hg*. We leveraged Ribo-seq to capture global trends of host and viral translation, to classify temporal gene sets, to identify differentially expressed processes, to detect co-regulated host and viral systems, and to gather evidence for post-transcriptional regulation in both the virus and its host. Our findings portray a model of the HFTV1 infection cycle and its effects on *Hg* (**Fig. 7**).

**Figure 7.**
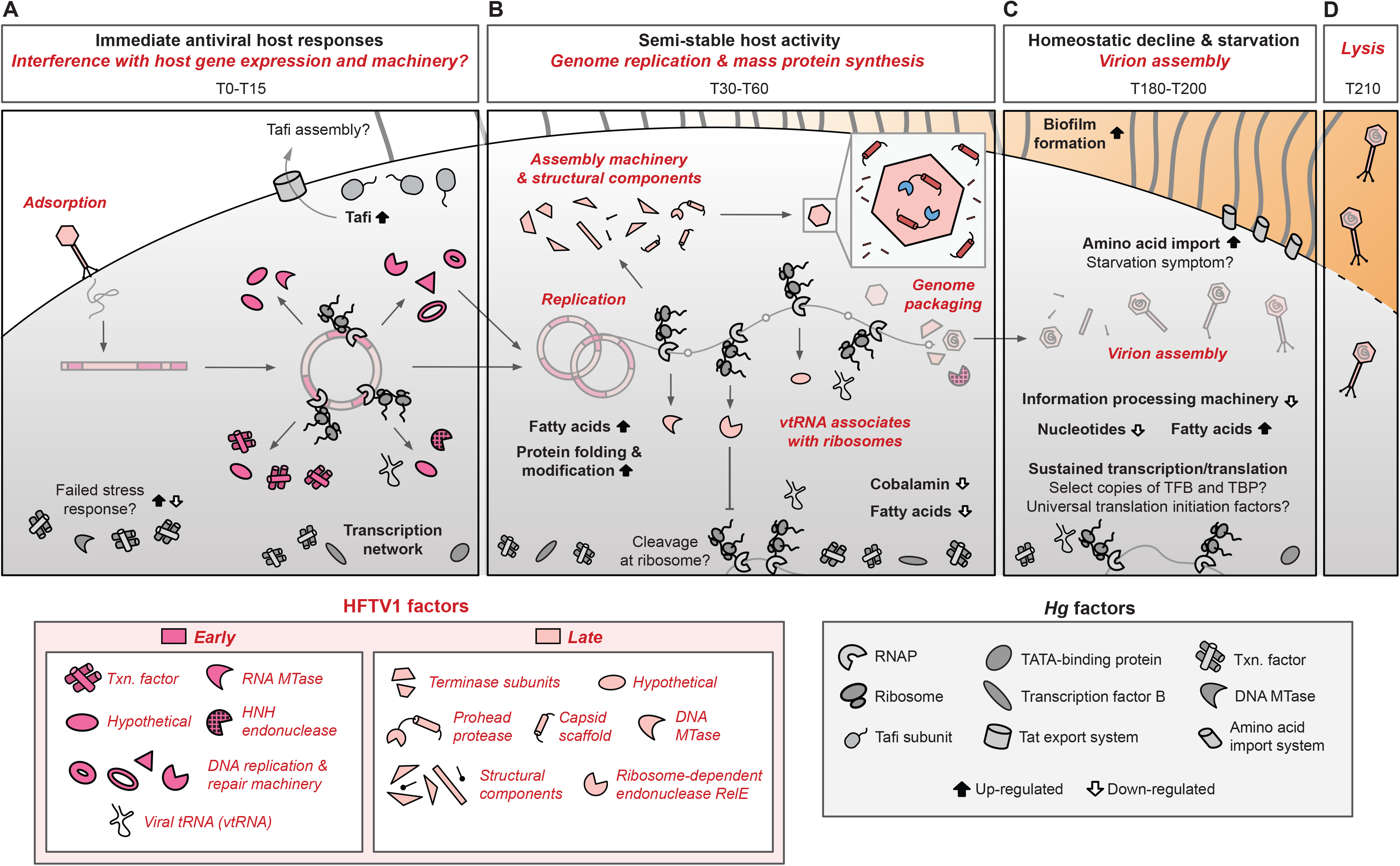
Model of the HFTV1 infection cycle. **(A)** Immediate host responses to viral infection, and potential interference by viral proteins with host gene expression and information processing machinery. **(B)** Semi-stable host activity, coinciding with viral genome replication and mass protein synthesis for progeny virions. **(C)** Homeostatic decline and starvation of the host during virion assembly. **(D)** Host lysis releasing progeny virions. Specific proteins are indicated in the key (*Hg* proteins in grey, early HFTV1 proteins in magenta, late HFTV1 proteins in pink). Up- and down-regulated *Hg* processes are respectively indicated with black upward- or white downward-facing arrows. All viral processes and factors are labeled in italics. Faded elements denote processes not addressed by this study but added for clarity; this includes ejection, circularization, replication, and packaging of the viral genome, as well as the assembly of the virion. Model is not to scale.

Regions of the viral genome were expressed rapidly after infection. This was evidenced at T0 both by the number of ribosome footprints associated with HFTV1 mRNA and by the modest co-purification of the HFTV1 vtRNA with host ribosomes. We suspect that the slightly shorter sample processing time for RNA-seq versus Ribo-seq accounts for the comparably low HFTV1 transcript abundance at T0. Early HFTV1 genes included multiple factors involved in DNA replication, establishing the viral genome as the dominant template for expression. They also included numerous transcription factors and hypothetical proteins, which may interfere with host gene expression or information processing machinery to facilitate viral gene expression or to inhibit antiviral defense systems.

The host responded quickly to infection in at least two ways: (i) there was a stress response involving transcription factors, a restriction-modification system, and universal stress proteins, although this response was quickly suppressed, perhaps by interference from early viral genes; (ii) there was induction of both a Tafi system and biofilm formation, the latter consistent with decreased motility previously observed during infection in *Hg* [58]. Since Tafi promote biofilm formation in other haloarchaea [96], we suggest that up-regulated Tafi were responsible for biofilm formation in *Hg* during infection. This may function as a defense mechanism similar to bacterial systems, in which biofilm inhibits phage diffusion and/or recognition of the host [103, 104]. Since *Hg* lacks a CRISPR-Cas system [57] and failed to mount a successful defense using its restriction-modification or toxin-antitoxin systems, biofilm formation may be one of its only means of defense. To our knowledge, these observations provided the first indications of biofilm formation as an antiviral defense in archaea.

In general, *Hg* exhibited a slow, gradual down-regulation of its information processing systems and associated metabolic activities through the infection cycle. Unlike other host-virus systems [105, 106], these changes were not rapid enough to suggest any global shutdown of transcription or translation, providing further evidence of the inability of *Hg* to counteract infection. Decreasing cobalamin biosynthesis may be an early indicator of decreasing nucleotide production. Overall, the gradual decline in host homeostasis likely facilitated viral activities such as genome replication and protein synthesis for virion assembly. Infection stressed the host and up-regulation of the thermosome and thioredoxins indicated that proteins, potentially from the host, had to be refolded and repaired. The most dramatic symptom of infection was the up-regulation of amino acid import systems immediately before lysis. This could indicate amino acid starvation caused by sustained viral protein synthesis, particularly for structural components of the virion.

A complex transcription network was active during infection, comprising both specific and basal transcription factors. Notably, several copies of the basal transcription factor TFB followed diverse expression patterns and co-clustered with other genes. The archaeal TFB is structurally and functionally similar to eukaryotic TFIIB but, like bacterial sigma factors, occurs in multiple copies expressed under specific conditions to enact widespread transcriptional regulation [107, 108]. In other haloarchaea, TFBs mediate responses to temperature shock [109, 110] and other environmental factors like light and oxygen [111]. Similarly, TFBs in *Hg* may be conditionally expressed during infection, coordinating the suites of genes with which they are co-clustered. This could make TFB—or select copies of it—a prime target for viral interference and efficiently reprogramming *Hg* during infection.

Throughout our analyses, we found evidence of post-transcriptional regulation in the genomes of both *Hg* and HFTV1. Certain host genes had high translation efficiency, including one for the pore that exports Tafi components (*tatA*) and several transcription factors from its aborted stress response. The expression of a subset of early HFTV1 genes also differed markedly between RNA-seq and Ribo-seq, including *gp07*, *gp46b*, and *gp62*, all encoding hypothetical proteins. These findings suggest that select genes in this archaeal host-virus system are post-transcriptionally regulated, some of which may play a crucial role in infection. Future work could explore the functions and regulation of these genes and their interplay with the mosaic nature of the archaeal information processing system.

One other HFTV1 gene clearly subject to post-transcriptional regulation was *gp17*, previously predicted to encode a single protein that performed both scaffolding and proteolytic functions. In contrast to its annotated function as a P2 gpO-like scaffolding/protease protein [87], *gp17* closely resembled “nested” protease-scaffold proteins, in which the full-length gene encodes the prohead protease, while an in-frame internal TSS encodes the scaffold protein.

This scheme appears in a few dsDNA viruses infecting bacteria and eukaryotes, including phage λ and Herpes simplex virus Type 1 (HSV-1) [87]. Although the proteases of phage λ, HSV-1, and HFTV1 showed weak pairwise sequence homology (20-23% ID), they shared predicted structural features, including a globular NTD and a disordered, helix-rich CTD (**Fig. S26**). The phage λ and HSV-1 proteases also have similar proteolytic activity, cleaving the scaffold, other structural components, and themselves. In HSV-1 specifically, the scaffold is degraded but discrete autoproteolytic fragments of the protease remain in the mature capsid [112]. The retention of fragments from across the protease—rather than just the CTD scaffold— parallels the roughly 1:1 ratio of the gp17 NTD and CTD in HFTV1 virions. We suggest that the full form of gp17 is the prohead protease and the truncated form is the capsid scaffold protein. As these proteins are distinguished by separate gene names in other viruses, we propose retaining *gp17* for the protease gene and naming its internal scaffold *gp17.5*, consistent with the nomenclature of HSV-1. These findings contribute to a model of the expression and function of gp17 and gp17.5 (**Fig. 3G**). Furthermore, the conservation of this transcript-level organization scheme lends a new form of support to the hypothesis that dsDNA head-tail viruses evolved from a common ancestor [15, 16, 30]. However, the mechanism underlying TSS selection may differ between viruses. In phage λ and HSV-1, the region directly upstream of the scaffold TSS has clear potential to form an RNA hairpin, whereas the equivalent region in HFTV1 does not (**Fig. S27**). Additionally, while phage λ harbors Shine-Dalgarno sequences upstream of both TSSs, we found no known motifs in HFTV1 that are associated with ribosome binding sites, consistent with the infrequent presence and use of Shine-Dalgarno sequences in *Haloferax* genomes [14]. The mechanism governing the protease-to-scaffold ratio therefore remains to be determined, though it seems independent of RNA secondary structure or ribosome binding motifs. Regulation by asRNA could also influence TSS selection, although we found no conclusive evidence of it in this study.

Viral translation may be partly mediated by the HFTV1 vtRNA, which is associated with host ribosomes during infection. Since the vtRNA is predicted to contain an intron, we suspect that our detection of two vtRNA species reflects the presence of a longer, unspliced variants and a shorter, spliced species; however, this and other processing or maturation steps remain to be verified experimentally. Surprisingly, while the vtRNA was predicted to decode the threonine codon ACG, there was no difference in ACG codon usage between the *Hg* and HFTV1 genomes. Consequently, we speculate that the vtRNA facilitates HFTV1 translation independent of codon usage, which could be clarified by assessing whether the HFTV1 vtRNA is aminoacylated, actively used during decoding, or used to preferentially translate specific viral transcripts. Further work is needed to define the maturation state, modification profile, and function of the HFTV1 vtRNA.

Together, our results depict HFTV1 infection as a finely orchestrated process in which *Hg* undergoes a gradual decline while the virus co-opts specific components of its information processing system. Canonical archaeal antiviral systems were either absent or failed in *Hg*, leaving biofilm production as the predominant defense mechanism, mediated by fimbriae co-regulated with early viral genes. Both host and viral genes showed evidence of post-transcriptional regulation. One example was *gp17*/*gp17.5*, which produced the prohead protease or the capsid scaffold protein depending on TSS selection, a mechanism found in other dsDNA viruses infection bacteria and eukaryotes. Host translation may be influenced by the HFTV1 vtRNA, which co-sedimented with host ribosomal complexes. By applying Ribo-seq paired with RNA-seq at high temporal resolution, our work captured global, *in vivo* transcriptional and translational dynamics during viral infection in an archaeon. This sets the groundwork for further study of post-transcriptional regulation, which was until now an unexplored layer of control in archaeal host-virus interactions.

## RESOURCE AVAILABILITY

RNA-seq reads, ribosome profiling reads, and annotation files are available on NCBI GEO: xxxxx (RNA-seq) and xxxxx (Ribo-seq). The mass spectrometry proteomics data have been deposited to the ProteomeXchange Consortium via the PRIDE (120) partner repository with the data set identifier xxxxx. Data analysis scripts are available on GitHub.

## ACKNOWLEDGEMENTS

We thank Michael Tassia and Rajiv McCoy for advice on our differential expression analysis pipeline; Neil Wood and Stephen Fried for mass spectrometry support for protocol design, analysis, and data interpretation; the HiLIFE Biocomplex Unit, University of Helsinki—a member of Instruct-ERIC Centre Finland, FINStruct, and Biocenter Finland— for providing access to ultracentrifugation services; Emma Dallon for sample collection; and Fijare Plous for pHGLR3 primer design. We are grateful to Hanna Oksanen for generously sharing HFTV1 and *H. gibbonsii* LR2-5. We thank Emma Dallon, Sadhana Chidambaran, and Isabel Baker for critical reading of the manuscript. This work was supported by the US National Science Foundation (grant no. MCB-2034271 to JDR) and Research Council of Finland (grant no. 354906 to LPS).

## SUPPLEMENTAL MATERIAL

**Figure S1.** Growth curves and 1-step growth curves for *Hg* infected with HFTV1 **(A)** Growth curves of *Hg* +/-HFTV1. **(B)** One-step growth curve of *Hg* infected with HFTV1. All experiments were performed at 42°C, 220 RPM, except for a viral adsorption period of 50 RPM, 5 min, immediately following infection or mock-infection.

**Figure S2.** Loss of *Hg* plasmid pHGLR3 in biological replicate #1 for RNA-seq/Ribo-seq. **(A)** Coverage by genetic element in each biological replicate of paired Ribo-seq (top) or RNA-seq (bottom). **(B)** Principal component analysis (PCA) of pHGLR3 Rlog-transformed gene expression data in biological replicates #2 and #3.

**Figure S3.** Growth and Ribo-seq QC for infected cells treated with HHT. **(A)** Growth of *Hg* with various doses of HHT. **(B)** Gene body coverage for HFTV1 (red) and Hg (grey) for cells treated with 1 mg/mL of HHT for 10 minutes. This method shows the ribosome density across the entire overall length (body) of each gene in contrast to a metagene analysis, which aligns genes at a single feature (e.g., start codon). Individual lines represent biological replicates. **(C)** Metagene analysis of +/-HHT Ribo-seq data for cells treated with 1 mg/mL of HHT for 10 minutes; the vertical lines on the graphs are positioned every 3 nt relative to 0 nt.

**Figure S4.** Reannotating the HFTV1 genome and mapping its translation start sites (TSSs) using HHT-treated cells. **(A)** HFTV1 reannotation pipeline with the number of genes processed at each stage. **(B)** Example of HHT enrichment data (normalized to -HHT data) along a segment of the HFTV1 genome. Peaks of HHT-induced ribosome density are visible at the 5’ ends of *gp21, gp22,* and *gp23*. Peaks such as these were used to identify and map TSSs. **(C)** Breakdown of HHT-mapped TSSs classified as canonical or non-canonical; non-canonical TSSs comprising those resulting in 5’ truncations or extensions of annotated ORFs, or the identification of internal or putative novel ORFs. **(C)** Example of 5’ extension of HFTV1 ORF *gp30* relative to reference annotation based on TSS mapping. **(D)** Example of the detection of a novel ORF (*gp52c*) and the removal of another (*gp52a*) based on TSS mapping. **(E)** Locations along the HFTV1 genome of revised features (extensions, truncations, internal ORF, and novel ORF / ORF removal) based on TSS mapping.

**Figure S5.** Read length distributions of Ribo-seq time course libraries across time points and biological replicates. "No data" indicates samples that were lost during ribosome isolation or library preparation

**Figure S6.** Codon frame analyses of Ribo-seq time course libraries across time points and biological replicates.

**Figure S7.** Metagene plots of Ribo-seq time course libraries across time points and biological replicates. Plots of HFTV1 genes are noisy likely due to the small number of coding sequences (∼80) used in the analysis; a similar level of noise is reproduced when metagene analysis is performed for *Hg* using a comparable number of coding sequences (not shown).

**Figure S8.** Sucrose gradient centrifugation polysome profiles of Hg cultures +/-HFTV1. **(A)** Hg cultures +HFTV1 collected from 30 to 180 minutes post-infection; 1.5 hr CTRL: uninfected control at 90 minute. **(B-C)** +/-HFTV1 ribosome fractionation from samples 60 minutes post-infection, undigested **(B)** or digested with MNase **(C)**.

**Figure S9.** Translation efficiency (TE) for *Hg* and HFTV1 genes across time points. **(A)** Depiction of all genes as points. **(B)** Depiction of only select genes of interest as points; same data as *Hg* genes in **(A)**. Black central marker shows distribution mean. Dashed lines are included at -1 and 1 on the log-transformed Y-axis, corresponding to raw TE values of -10 or 10, respectively. TE was calculated per gene in the following manner: TE = (*Rlog SFN* Ribo-seq read counts + 1)/(*Rlog SFN* RNA-seq read counts + 1). TE values were log-transformed for these graphs for visualization purposes. *Rlog:* regularized log-transformed (*DESeq2*). *SFN: sizeFactor-*normalized (*DESeq2*).

**Figure S10.** Ribo-seq clusters for differentially expressed genes over the time course. **(A)** Scaled *Rlog SFN* reads. **(B)** L2FC relative to T0; shaded area covers L2FC +/-1.5. HFTV1 genes: red. *Hg* genes: black. *Hg* plasmid pHGLR3 is not represented (see Method Details). *Rlog:* regularized log-transformed (*DESeq2*). *SFN: sizeFactor-*normalized (*DESeq2*). L2FC: Log2 fold-change.

**Figure S11.** Ribo-seq clusters for differentially expressed genes on *Hg* plasmid pHGLR3 over the time course. **(A)** Scaled *Rlog SFN* reads. **(B)** L2FC relative to T0; shaded area covers L2FC +/-1.5. *Rlog:* regularized log-transformed (*DESeq2*). *SFN: sizeFactor-*normalized (*DESeq2*). L2FC: Log2 fold-change.

**Figure S12.** RNA-seq clusters for differentially expressed genes over the time course, shown as **(A)** scaled *Rlog SFN* reads or **(B)** L2FC relative to T0; shaded area covers L2FC +/-1.5. HFTV1 genes: red. *Hg* genes: black. *Hg* plasmid pHGLR3 is not represented (see Method Details). *Rlog:* regularized log-transformed (*DESeq2*). *SFN: sizeFactor-*normalized (*DESeq2*). L2FC: Log2 fold-change.

**Figure S13.** RNA-seq clusters for differentially expressed genes on *Hg* plasmid pHGLR3 over the time course, shown as scaled *Rlog SFN* reads (top) or L2FC relative to T0; shaded area covers L2FC +/-1.5. *Rlog:* regularized log-transformed (*DESeq2*). *SFN: sizeFactor-*normalized (*DESeq2*). L2FC: Log2 fold-change.

**Figure S14.** Ribo-seq cluster families for differentially expressed genes over the time course, shown as scaled, Rlog-transformed *SFN* reads (top) or L2FC relative to T0 (bottom). Gene clusters derived from *Hg* plasmid pHGLR3 could not be processed at the same time as other clusters due to missing data (see Method Details), so they were assigned manually to the cluster families produced by clusters derived from all other genetic elements.

**Figure S15.** Extended data of Ribo-seq cluster families. This is a combination of **Fig. 2**, **Fig. 5**, and additional data to provide a comprehensive view of Ribo-seq cluster families and the functions of their associated genes. **(A)** Analysis pipeline to identify and cluster dRibo genes. Only genes that corresponded to CDSs were used for differential expression analysis. **(B)** Cluster families produced by clustering mean expression trajectories of gene clusters; visualized using scaled *Rlog SFN* reads. **(C)** Numbers and proportions of *Hg* and HFTV1 genes in each cluster family.CDS: Coding sequence. **(D)** Abundance of functional categories for HFTV1 genes. **(E)** Proportions of dRibo and non-dRibo *Hg* genes per arCOG functional category. **(F)** Enrichment analysis of functional categories in each cluster family restricted to *Hg* genes. (*): adjusted p-value < 0.05. *Rlog:* regularized log-transformed (*DESeq2*). *SFN: sizeFactor-* normalized (*DESeq2*).

**Figure S16.** Additional expression patterns of *gp17*. **(A)** Translation efficiency of *gp17* over the course of infection. Translation efficiency was calculated per time point as follows: Translation efficiency = (Ribo-seq *SFN* reads + 1)/(RNA-seq *SFN* reads + 1). **(B)** Genome viewer representation for the region of the HFTV1 genome containing *gp17* to *gp39*. **(C)** Magnification of region from *gp17* to the 5’ end of *gp19*. Note the difference of scale of antisense coverage tracks required to visualize patterns in **(C)** compared to **(B)**. (+)-strand features colored in red. (-)-strand features colored in dark red / maroon. All RNA-seq coverage data are from T180.

**Figure S17.** Quality control metrics of LC-MS/MS data on infected cells (T30 and T180, three biological replicates each) or purified virions (three technical replicates, split immediately prior to reduction with DTT). **(A)** Cumulative protein abundance for *Hg* or HFTV1 for T30, T180, and purified virions. **(B)** Number of unique proteins detected for *Hg* or HFTV1 for T30, T180, and purified virions. **(C)** Rank abundance curves for each sample; individual proteins colored in black for r *Hg* and red for HFTV1. **(D)** Same as **(C)**, but showing only HFTV1 proteins. **(E)** Volcano plot of differential abundance of HFTV1 proteins in purified virions versus in cells at 180 min post-infection (T180). All structural components are enriched in purified virions.

**Figure S18.** SDS-PAGE of samples collected during PEG-free HFTV1 purification process. Clarified lysate: *Hg* lysate after pelleting cell debris. Filtered lysate: clarified lysate after 0.2 um filtration. Pellet I supernatant: supernatant after ultracentrifugation of filtered lysate. Pellet II supernatant: supernatant after ultracentrifugation of sucrose gradient light-scattering zones (expected to comprise full virions), diluted in 18% salt water. Pellet II resuspension (1x): final purified virus sample used for further processing for mass spectrometry.

**Figure S19.** Proportions of dRibo and non-dRibo genes in RNA-seq and Ribo-seq.

**Figure S20.** Ribo-seq expression patterns of *Hg* translation initiation factors. Expression patterns as L2FC hierarchically clustered solely for visualization purposes using *pheatmap*. N/A for "Cluster family" indicates that the corresponding gene did not exhibit differential ribosome density. N/A for "Bacterial homolog" indicates that the corresponding archaeal product has no bacterial homolog.

**Figure S21.** T200 vs. T150 time point analysis of Ribo-seq genes differentially expressed. **(A)** *Hg* genes. **(B)** Proportion of all differentially up-regulated *Hg* genes (corresponding to region of **(A)** in pink box) that encode ABC-type oligopeptide import components. **(C)** HFTV1 genes at the same y-axis scale as **(A)** (left) or with magnified y-axis scale to visualize marginally down-regulated genes (right). Thresholds for differential expression: |L2FC| > 1.5, adjusted p-value < 0.05.

**Figure S22.** Signal peptides of Tafi pilin homologs. Predictions performed with SignalP.

**Figure S23.** Signal peptides and predicted structures of putative *Hg* Tafi tip TafC2 and similar proteins. **(A-B)** *Hg* TafC2. **(C-D)** *Natrinema* TafF2. **(E-F)** *E. coli* AG43. **(G)** Phage P22 tailspike protein; no signal peptide present, presumably since this product does not require export across a membrane. For all proteins, signal peptides predicted with SignalP and structures predicted by AlphaFold2 (Rank 1 models shown). Signal peptides on structures indicated in light green with present.

**Figure S24.** Multimeric predictions for *Hg* Tafi structural components (products of 13635, 13645, and 13655). **(A-B)** Dimeric interactions, either **(A)** heterodimeric or **(B)** homodimeric. **(C)** Trimeric interaction. All multimeric predictions performed with AlphaFold2 (Rank 1 models shown).

**Figure S25.** Quantitative RT-PCR (RT-qPCR) of *Hg* TafA +/-HFTV1. All analyses were conducted on T30 samples from the same biological replicates used for RNA-seq and Ribo-seq. Reference gene *gyrB* (locus tag HfgLR_RS07905) selected based on minimal change in expression at T30 in RNA-seq data (T30 vs. T0 L2FC = 0.043). Primer efficiencies were > 90%. **(A)** RT-qPCR amplification curves. **(B)** L2FC of *tafA* based on ΔΔCT calculation, using *gyrB* as reference gene, *tafA* as target gene, +HFTV1 as experimental condition, and -HFTV1 as control condition, 5 technical replicates per dilution per gene, 3 biological replicates.

**Figure S26.** Structural comparisons of proteases and scaffold proteins from phage lambda, HSV-1, and HFTV1. Structures predicted by AF2; Rank 1 structures shown.

**Figure S27.** Features of protease/scaffold sequences in dsDNA head-tail viruses. Homology, functions, and regulation of nested dsDNA protease/scaffold proteins informed by [87, 112, 114–117].

**Table S1.** Read counts of Ribo-seq libraries throughout processing. **Table S2.** Read counts of RNA-seq libraries throughout processing. **Table S3.** Documentation of translation start site mapping.

**Table S4.** Classification of genes as dRibo or non-dRibo.

**Table S5.** Classification of genes as dRNA or non-dRNA.

**Table S6.** Master spreadsheet of genes, including cluster family assignment, cluster assignment, functional annotations, and various expression values.

**Table S7.** Protein abundance for infected cells and purified virions by LC-MS/MS.

**Table S8.** Differential protein abundance results for purified virions vs. HFTV1-infected *Hg* 180 minutes post-infection.

**Table S9.** Primers used in this study (Ribo-seq library preparation; RT-PCR; pHGLR3 presence/absence; RT-qPCR; Northern blot probes).

**Table S10.** Summary of major commands and tools for Ribo-seq and RNA-seq data processing.

## STAR METHODS

### EXPERIMENTAL MODEL AND STUDY PARTICIPANT DETAILS

*Haloferax gibbonsii* LR2-5 [24, 57] (*Hg*; genome assembly ASM1496974v1, DSM 112399) and HFTV1 virus [24] (DSM 117943).

### METHOD DETAILS

#### Cell growth

*Hg* was grown at 42 °C, 220 RPM (Eppendorf Innova S44i), in liquid modified growth medium (MGM) [118] consisting of 18% buffered salt water (BSW; 2.46 M NaCl, 88.8 mM MgCl2.6H2O, 85.2 mM MgSO4.7H2O, 56.4 mM KCl, 3 mM Tris-HCl pH 7.5) [118], yeast extract (Fisher BP1422-500), and peptone (Oxoid LP0037B).

#### Viral infection

*Hg* was grown to stationary phase (OD_600_ > 1), back-diluted to OD_600_ = 0.01 in fresh medium, regrown, and infected with HFTV1 during mid-exponential phase (OD_600_ ∼0.6) with a multiplicity of infection (MOI; approximate number of virions per cell) of 10; all infection experiments reported in this study were performed using purified HFTV1 (described below). Viral adsorption was facilitated by decreasing shaking speed to 50 RPM for 5 min immediately after infection, after which shaking speed was returned to 220 RPM. All infections in liquid culture were performed in flasks. Growth on agar plates was performed on solid MGM. Under these conditions, we found that lysis began at ∼3.5-4 hours post-infection (**Fig. S1**).

#### Viral purification

For HFTV1 purification, *Hg* culture was infected as described above, then moved to +37 °C and incubated for 36-48 h with shaking decreased to 80 rpm. After complete lysis (OD_600_ <0.2), the lysate was clarified by centrifugation at 10,800*×g*, 30 min at +4 °C. To precipitate HFTV1, clarified lysate was mixed with PEG6000 to final concentration at 4% and incubated stirring at +4 °C for 1 h. PEG-precipitated virus was collected by centrifugation at 10,800×*g*, 40 min, +4 °C and pellet was dissolved in 18% SW buffer [118] and incubated O/N at +4 °C. The undissolved particles were removed by centrifugation at 6,300×*g*, 10 min at +4 °C. The viruses were purified by rate zonal centrifugation in 20-50% sucrose gradient using a AH-629 rotor (Sorvall) at 103,586×*g* for 3h at +15 °C. The virus containing zones were collected, diluted with 18% SW buffer in 1:2-1:3 ratio (virus zone:buffer) and virus particles were pelleted by centrifugation in a T-647.5 rotor (Sorvall) at 113,580×*g*, 2 h, +4 °C. The final virus pellet was resuspended in 18% SW buffer and stored at -70 °C. The infectivity and protein concentration were determined by plaque and Bradford assays, respectively. Viral stock purified by this PEG-based method was used for all experiments described in this paper except for proteomic analysis of purified viral particles. Due to the interference of PEG with mass spectrometry [119], HFTV1 particles used for proteomic analysis were purified independent of PEG6000. First, cultures were infected and lysate was clarified in the manner described above for PEG-based purification. After clarification, lysate was filtered through a 0.2 µm PES filter (Millipore S2GPU05RE) by gentle vacuum. Viral particles were pelleted from filtered lysate by ultracentrifugation using a 45Ti rotor (Beckman) at 113,580x*g*, 2.0 hours, +15 °C. Supernatant was removed by decanting. Pellets were washed gently with 18% SW buffer and initially resuspended by pipetting, then further dissolved and resuspended overnight in 18% SW buffer with agitation. Resuspended viral particles were isolated along a 10-40% sucrose gradient by ultracentrifugation using a SW41Ti (Beckman) rotor at 30,000 RPM (154,049x*g*), 2 h, +4 °C. Gradient zones containing the fully intact virus were then isolated and further processed in the same manner described for PEG-based stocks. Aside from infections to assess stock infectivity and quality, viral stock purified by this PEG-free method was used only as input material for mass spectrometry.

#### Genome-based revisions of the HFTV1 annotation

The HFTV1 annotation was retrieved from NCBI (genome assembly ASM420877v1, RefSeq assembly GFC_004208775.1). Additional features in the HFTV1 genome were predicted *in silico* using the wrapper program Pharokka v. 1.7.5 [77, 120]. Within Pharokka, coding sequences (CDSs) were predicted using PHANOTATE [121], tRNAs were predicted using tRNAscan-SE 2.0 [122], tmRNAs were predicted using Aragorn [123], and CRISPRs were predicted using CRT [124]. Functional annotation was performed by comparing each CDS to the PHROGs [125], VFDB [126] and CARD [127] databases using MMseqs2 [128] and PyHMMER [129]. All Pharokka-predicted features were further revised using Phold [120, 130]. Within Phold, all protein sequences were translated to the 3Di token alphabet using ProstT5 [131] and then searched against the Phold phage protein database using Foldseek [132]; the Phold database comprises over 1.36 million phage protein structures primarily predicted by Colabfold [133].

Feature descriptions were also updated based on the recent cryo-EM structure of HFTV1 virions [28]. Unique features predicted by the Pharokka/Phold pipeline relative to the HFTV1 NCBI annotation were compared to a previously published revision of the HFTV1 annotation [58]; a detailed form of this annotation was not made available with the previous publication, so comparisons were performed qualitatively with the published figures. A revised HFTV1 annotation file from our work will be made available upon publication.

#### Screening for loss of native *Hg* plasmid pHGLR3

One biological replicate of paired RNA-seq/Ribo-seq data showed extremely low coverage of the native *Hg* plasmid pHGLR3 (**Fig. S2**), consistent with spontaneous plasmid loss previously reported in *Hg* [134]. Given the consistent trends in host and viral reads and in viral gene expression across all other biological replicates, we concluded that pHGLR3 genes do not play a decisive role in the HFTV1 infection. Following this observation, we screened all *Hg* colonies by PCR and restricted major experiments to pHGLR3-positive colonies. To accommodate pHGLR3 absence in one replicate, pHGLR3 data were analyzed separately in PCA and downstream differential expression analyses (**Fig. S2**). *Hg* colonies were screened by PCR (NEB Phusion polymerase, M0530S) with pHGLR3-specific primers (**Table S9**).

#### Ribo-seq sample collection

Ribo-seq samples were collected by growing and infecting cultures as described above, allowing infection to progress to the desired time point, and then flash-freezing culture as previously described (Gelsinger et al., 2022) by dispensing it dropwise into liquid nitrogen; this produced small frozen “beads” or “dippin’ dots” of culture that were later processed for ribosome fractionation and library preparation.

#### Ribo-seq and RNA-seq infection time course

RNA-seq and Ribo-seq samples at 0, 5, 15, 30, 60, and 180 min post-infection were collected from identical biological replicates (only 2-3x biological replicates of Ribo-seq were sequenced per time point due to sample loss during library preparation). At each time, 20 mL of culture were pelleted at 6,000xg, +4 °C, 5 min. The supernatant was removed and the pellets were snap-frozen in liquid nitrogen; these cell pellets were later processed for RNA-seq.

Simultaneously, the remaining volume (100 mL) of infected culture was flash-frozen as described in "Ribo-seq sample collection" above. In parallel, the 6x -HFTV1 RNA-seq cultures (mock-infection) and the -HFTV1 growth control culture were also collected. The time at which cultures were inoculated with HFTV1 or mock-infection medium was treated as 0 min post-infection, before the 5 min adsorption period at 50 rpm described in "Viral infection" above. For Ribo-seq used to assess late stages of infection, 125 mL infected cultures were incubated until 200 minutes post-infection, after which they were harvested as described in "Ribo-seq sample collection" above (2x biological replicates).

#### Ribo-seq for translation start site mapping

To determine an appropriate dosage of harringtonine (HHT) to arrest translation in *Hg* for translation start site (TSS) mapping, starter cultures of *Hg* were grown to stationary phase, then back-diluted to OD_600_ = 0.1 in 18% MGM containing HHT (Sigma-Aldrich, #SML1091) dissolved in DMSO with final dosage of 0.1, 0.2, 0.5, or 1 mg/mL. Control cultures were spiked with DMSO or MGM at a volume equivalent to the maximum HHT dosage. Culture growth was measured from 0 to 30 hours after back-dilution using a multi-well Bioscreen plate (Bioscreen C Pro), minimum 3x technical replicates per condition. This experiment was performed for one biological replicate due to the inhibitory cost of HHT when used at a dosage of 1 mg/mL, even for small culture volumes.

For Ribo-seq used to map HFTV1 TSSs, 125 mL infected cultures were incubated until 150 minutes post-infection, then treated with either HHT at 1 mg/mL (+HHT samples) or DMSO (- HHT samples) for 10 min before harvesting the cells as described above (3x biological replicates).

### Ribo-seq read processing and quality control

Only forward reads were processed for Ribo-seq analysis. Adapters were trimmed using Trim Galore (v. 0.6.10) [135]. Reads were aligned to rRNA using Bowtie (v. 1.3.1) [136]. Reads that did not align to rRNA (i.e., rRNA-filtered reads) were aligned to a merged *Hg*-HFTV1 genome file using Bowtie (v. 1.3.1) [136]. The merged genome file was generated from the *Hg* genome assembly ASM1496974v1 and the HFTV1 genome assembly ASM420877v1; this file will be made available upon publication. Alignment files were generated, sorted, and indexed using Samtools (v.1.20) [137]. Coverage files were generated with deepTools *bamCoverage* [138]. For differential expression analyses, reads per feature were calculated in a strand-specific manner using featureCounts [139] for a merged *Hg*-HFTV1 annotation file constructed from the *Hg* annotation accompanying the NCBI RefSeq assembly GCF_014969745.1 and the revised HFTV1 annotation described above. Reads that overlapped more than one feature were counted as a fraction of a read, with the denominator determined by the number of overlapped features. Specific commands and parameters are listed in **Table S10**. Various quality metrics of raw reads, trimmed reads, and aligned reads were assessed using FastQC [140]. Other statistics for genome-aligned reads were calculated using samtools (*stats* and *coverage*; v. 1.20) [137]. Gene body coverage of aligned reads was assessed using RSeQC *geneBody_coverage* (v. 5.0.4) [141, 142]. Metagene analysis was performed using *plastid* (*metagene* module) [143]. Briefly, reads were subset to those mapping to CDSs, aligned at the start codon of each CDS, and then used to calculate ribosome density based on the mean number of 3’ read ends at each nucleotide position 250 nt into the CDS. An additional 50 nt upstream of the start codon was also included in the analysis to compare ribosome density within and outside of CDSs. The distance between the P-site and the 5’ end of the ribosome footprint (i.e., the P-site offset) was determined using *plastid* (*psite* module) [143]. This analysis was similar to metagene analysis (described above), but it stratified ribosome density by read length; a consistent offset between a start or stop codon and a spike in high ribosome density (due to the rate-limiting nature of translation initiation and termination) should be evident across various read lengths. For the current work, the P-site offset was measured as the distance between the stop codon and a spike of high ribosome density shortly upstream of the stop codon. The start codon was forgone since *Hg* has a high proportion of leaderless transcripts, resulting in high ribosome density precisely at the start codon and no measurable P-site offset for this feature; we previously used a similar approach in *Haloferax volcanii* [79].The P-site offset was found to be 14 nt for abundant ribosome footprint lengths (24 and 27 nt) associated with elongation. The proportion of ribosome footprint ends mapping to each position within a codon (0, 1, or 2) was calculated using *plastid* (*phase_by_size* module) [143]. For this quality assessment, we used ribosome footprints that were 27 nt in length and a P-site offset of 14 nt *relative to the 5’ end* of each read. This method is consistent with our previous analysis in *Haloferax volcanii* [79].

### Revision of 5’ ORF boundaries by translation start site mapping with +/-HHT Ribo-seq

We revised the 5’ boundaries of HFTV1 ORFs by mapping translation start sites (TSSs) in the viral genome based on Ribo-seq data +/-HHT. For this application, the P-site was determined differently compared to libraries used solely for differential expression analysis. Based on prior quality assessments, our Ribo-seq reads across all libraries exhibited greater variation in length at their 5’ ends compared to their 3’ ends. This did not cause issues for our quality assessments of libraries for differential expression analyses because the downstream codon frame analysis (the only application of the P-site offset in those libraries) was performed only on reads of a single length (27 nt). However, for TSS mapping, we sought to retain as many reads as possible without subsetting the data by length. Additionally, 5’-mapping in metagene analyses revealed a sharp peak in ribosome density at the 0 nt position relative to the CDS. This was previously observed in other Ribo-seq datasets [144, 145] and is likely attributable to a cloning bias from the well-defined 5’ end of transcripts; this bias is revealed when similar sequences such as start codons are aligned. Consequently, due to i) the variable length of reads at their 5’ ends and ii) the artifactual 5’-based enrichment of ribosome density at the 0 nt position, we determined the P-site offset relative to the 3’ ends of reads to resolve TSSs. This was achieved by comparison of metagene plots generated with *plastid* (*metagene* module) [143] for libraries +/-HHT. The highest peak downstream of the start codon, elevated in the +HHT libraries relative to the -HHT libraries, and upstream of the elongation-associated region of 3 nt periodicity was taken as the indicator of the 3’ boundary of initiating ribosomes (**Fig. S3**). By this approach, we identified a P-site offset of 15 nt *relative to the 3’ end* of each read. After the selection of a P-site offset, all genome-aligned reads from Ribo-seq libraries +/-HHT (150 minutes post-infection) were trimmed to comprise only the nucleotide at the start of the ribosomal P-site using deepTools *bamCoverage* [138]. Coverage of the resulting 1-mers across the merged *Hg-*HFTV1 genome was adjusted for library size by calculating bins per million with a bin size of 1 (i.e., equivalent to reads per million (RPM)) using deepTools *bamCoverage*. Coverage data were then averaged across three biological replicates using deeptools *bigwigAverage*. In the same step, we normalized treated data by untreated data and log-transformed in the following manner log2((+HHT RPMs + 1)/(-HHT RPMs + 1). All of these steps were performed in a strand-specific manner to ultimately produce coverage files for the forward and reverse strand of the HFTV1 genome. Values in these files corresponded to enrichment or depletion of ribosome density induced by HHT treatment relative to untreated controls; we refer to these data hereafter as HHT enrichment data. The HHT enrichment data were scanned manually in IGV (ref, version). Data for the reverse strand were found at this stage to be very noisy, so further analysis was conducted only on the forward strand. We searched for 5’ extensions and truncations based on the appearance of HHT-enriched ribosome density in positions that differed relative to the nearest annotated TSS in the genome. We also searched for novel ORFs based on the appearance of HHT-enriched density in positions that had no explanatory ORF on the forward strand. In total, 9 revisions (5x 5’ extensions, 3x 5’ truncations, and 1x novel ORFs) were incorporated into the HFTV1 annotation. This reannotation, along with the features derived from Pharokka/Phold, was then used for our differential expression analysis and will be made available upon publication. The unusual features of *gp17* and its domain-specific expression patterns were excluded from this annotation and were explored with separate analyses.

### Ribo-seq sample processing, library preparation, and sequencing

Flash-frozen liquid culture was processed further for Ribo-seq as previously described (Gelsinger et al., 2022). Briefly, batches of 50 g of frozen culture were supplemented with 100 µg/mL anisomycin (Sigma-Aldrich, #A9789) and pulverized in a cryomill (Spex, 6870 Large Freezer/Mill) for 8× cycles (1 min grinding at 10 Hz, 1 min cooling). Cell lysates were thawed at room temperature and clarified at 13,000×g, 15 min, 4 °C. Clarified lysate was loaded onto a 1 M sucrose cushion (sucrose dissolved in 1× lysis buffer [3.4 M KCl, 100 mM MgCl_2_, 50 mM CaCl_2_, 10 mM Tris-HCl pH 7.5]) and ribosomes were pelleted using a 45Ti rotor (Beckman Coulter), 40,000 RPM (185,511xg), 3 h, 4 C. Ribosome pellets were washed once with 1x lysis buffer, then resuspended in 300 µL 1x lysis buffer. RNA concentrations of resuspended pellets were measured using the Qubit RNA Broad Range assay (Thermo Fisher Scientific, #Q10210). Unprotected mRNA was then digested by adding micrococcal nuclease A (MNase A) to the ribosome pellets for 1 h, 25 °C, at a ratio of 350 U MNase A (purified in the lab as described in [79] per 1 µg RNA. MNase-treated ribosome resuspensions were then loaded onto a 10-50% sucrose gradient (sucrose dissolved in 1x lysis buffer) and their components were separated by ultracentrifugation with a SW41Ti rotor (Beckman Coulter), 273,620x*g*, 2.5 h, 4 °C. Fractions were collected along each gradient in 450 µL volumes while measuring A260 using a BioComp Piston Gradient Fractionator, flash-frozen on dry ice, and stored at –80 C until further processing. TRIzol LS (Invitrogen, 10-296-010) was used per manufacturer instructions to extract RNA from the sucrose gradient fractions that corresponded to 70S monosomes. The resulting RNA was separated by size using a 15% polyacrylamide TBE-urea gel. RNA fragments in the range of 10-45 nt were excised, eluted from gel fragments, precipitated by isopropanol, treated with T4 polynucleotide kinase (NEB, #M0201S), ligated to a universal linker (NEB, Universal miRNA Cloning Linker, #S13115S) using T4 RNA ligase (NEB, #M0242), purified (Zymo, Oligo Clean and Concentrator Kit, #D4061), and reverse-transcribed (Invitrogen, SuperScript III, #18080044) using custom primers (Gelsinger et al., 2022). Resulting cDNA fragments were purified on a 10% polyacrylamide TBE-urea gel, circularized (EpiCentre, CircLigase, CL4115K), and amplified by PCR (NEB, Phusion polymerase, M0530S) for 8-12x cycles using custom primers (**Table S9**) [144]. The optimal number of cycles was selected for each library by analysis of pilot PCR products on an 8% polyacrylamide TBE gel. Final library material was produced using the optimal number of cycles for each library. The amplified cDNA was separated on an 8% polyacrylamide TBE gel, excised at the expected size range, eluted from gel fragments, and precipitated by isopropanol. Resuspended cDNA libraries were assessed for purity, size, and concentration by chip-based electrophoresis (Agilent BioAnalyzer, High-Sensitivity DNA Kit, 5067-4626). Libraries were sequenced by Novogene (150 bp paired-end reads, NovaSeq).

### RNA-seq sample processing, library preparation, and sequencing

RNA was extracted from cell pellets using a Zymo Quick-RNA Miniprep Kit (R1054), treated with DNase I (NEB, M0303S) at 37 °C, 15 min, and cleaned and concentrated using a Zymo RNA Clean & Concentrate Kit (R1017). RNA concentration was measured using a Qubit RNA Broad Range Assay kit (Invitrogen, Q10210). Total RNA was assessed for integrity by chip-based electrophoresis (Agilent, Bioanalyzer RNA 6000 Pico Kit, 5067-1513). rRNA depletion using custom probes for *Hg,* library preparation using Illumina’s Stranded Total RNA Prep Ligation with Ribo-Zero Plus kit and 10 bp unique dual indices, and sequencing to produce 150 bp paired-end reads using the Illumina NovaSeq X Plus platform were performed by SeqCenter (Pittsburgh, PA, USA).

### RNA-seq read processing and quality control

RNA-seq paired-end reads were processed as paired, in contrast to Ribo-seq reads. Otherwise, RNA-seq reads were handled in the same manner described for Ribo-seq (including: adapter trimming; rRNA filtration; alignment to a merged *Hg*-HFTV1 genome; generation of alignment, coverage, and statistics files; calculation of reads per feature; and quality assessment). Specific commands and parameters are listed in **Table S10**.

### Homology and motif analyses

Gene and protein sequences were assessed for homology using BLAST protein alignment [146]. Protein domains were scanned for conserved domains using InterPro [147] as hosted on EMBL-EBI (https://www.ebi.ac.uk/). Signal peptides were detected using SignalP v. 6.0 [97], both hosted through DTU ( https://services.healthtech.dtu.dk/services/SignalP-6.0/) and as a component of InterPro. Protein structures were predicted using ColabFold [133], which relies on AlphaFold2 [148] and MMSeqs [128]. Protein structures were visualized using Mol* Viewer [149] and assessed for homology using FoldSeek [132]. Local protein alignments performed with EMBOSS Water hosted on EMBL-EBI [150].

### Relative synonymous codon usage calculation

The HFTV1 and *Hg* coding sequences were extracted from their respective genomes using a custom R script. Start and stop codons were removed. RSCU values were calculated and visualized by RSCUcaller [151].

### Prediction of vtRNA structures

Both structures of vtRNA, with and without intron, were predicted in Snapgene v. 8.2.2 using RNAfold and RNAsubopt from ViennaRNA package [89]. For the prediction, default parameters and a temperature of 42°C were used.

### Northern blot analysis of vtRNA expression during HFTV1 infection

The detection of vtRNA and host tRNA expression during infection time course was performed on the same RNA samples used for RT-PCR. 3 µg of total RNA were separated on 10% PAA/7M urea denaturing gels and (v)tRNAs were detected by chemiluminescent Northern blot as previously described [152], using the probes listed in **Table S9.** For detection of vtRNA association with translating ribosomes, polysome profiles at 0 and 180 min p.i. were prepared as described for Ribo-seq but without the nuclease digestion step. Total RNA was isolated from the polysome fractions using TRIzol LS (Invitrogen, 10296010) as described above. Fractions were pooled according to their sedimentation position into free-lysate, small subunit (30S), large subunit (50S), monosome (70S), polysomes (light and heavy) and remaining large complexes. The detection of vtRNA and host tRNA expression across polysome profiles was performed as described above

### RT-PCR and RT-qPCR

For RT-PCR of select early and late HFTV1 genes, the overnight Hg culture (<40 h) was diluted to OD_600_ ∼0.1 in fresh 18% MGM and grown at 42 °C, 200 rpm until OD600 = 0.6. The 14 parallel 20 mL cultures were infected with either purified HFTV1 (MOI = 10) or mock-infected with 18% SW buffer and after HFTV1 adsorption (50 rpm, 5 min) cultures were grown further at 42 °C, 200 rpm. Individual cultures (+HFTV1 and -HFTV1) were collected by centrifugation at 8000 g for 5 min at time points 0, 5, 15, 30, 60, 120 and 180 min p.i. Pellets were resuspended in 5 mL Trizol (38% acidic phenol, 0.8M guanidine thiocyanate, 0.4M ammonium thiocyanate, 0.1M sodium acetate, pH 5.3, 5% glycerol) (ref), flash frozen in liquid nitrogen and stored at - 70 °C until further processed. To isolate total RNA, the pellets in Trizol were thawed at RT and 1 mL of 1-bromo-3-chloropropane was added. Samples were vigorously vortexed (1-2 min) and then incubated for 10 min at RT. Phases were separated by centrifugation at 10,000×g, 15 min at RT. Aqueous phase was collected to new tube and precipitated by addition of 2.5 mL of 99.6% ethanol. RNA precipitation was carried out O/N at -20 °C. Next, the RNA was pelleted by centrifugation at 10,000×g, 45 min, +4 °C. Supernatant was removed, and RNA pellet was washed by 1 mL of 80% EtOH. Total RNA pellets were air-dried and dissolved in RNase/DNase-free water. 50 μg of total RNA were treated by 20 U of RQ1 DNase (Promega, M6101) according to the manufacturer’s instructions, followed by RNA re-extraction by acidic phenol/BCP. cDNA was prepared by reverse transcription of 1 μg DNased total RNA and 100 pmol random hexamers (Qiagen, 79236) using Maxima RT (ThermoFisher Scientific, EP0741) according to the manufacturer’s instructions. PCR reactions consisted of 0.25 μM (each) forward and reverse primers, 1 μL undiluted cDNA and 1x HOT FIREPol mix (SolisBiodyne, 04-36-00120) and were run for 20 cycles. RT-PCR products were analyzed on 2% agarose gels.

Primers are listed in **Table S9.**

Reverse-transcription quantitative PCR (RT-qPCR) was performed on the same RNA that was extracted for RNA-seq and paired with Ribo-seq. For three biological replicates at 30 min post-infection under both infected (+HFTV1) and mock-infected (-HFTV1) conditions, DNase-treated RNA was reverse-transcribed into cDNA using the SuperScript III First-Strand Synthesis system with random hexamers (Invitrogen, 18080051); control reactions with RNA but no reverse transcriptase (-RT) were included for all samples. All qPCR steps were performed in 96-well plates using PowerUp SYBR Green Master Mix (Applied Biosystems, A25742) with the BioRad C1000 Thermal Cycler and CFX96 Real-Time System.

### Mass spectrometry sample preparation

Mass spectrometry was performed on HFTV1-infected *Hg* (3x biological replicates) as well as PEG-free purified HFTV1 virions. At 30 min and 180 min post-infection, culture volumes of 20 mL per time point were pelleted at 8,000xg, 5 min, 4 °C. Supernatant was removed, then pellets were flash-frozen and stored at -80 °C. Pellets were later washed with phosphate-buffered saline (PBS) and resuspended in 1 mL denaturing buffer (8 M urea, 50 mM AmBic, pH 8, freshly prepared). Cells were lysed and protein was solubilized by sonication on ice (VWR Symphony) for 6x cycles (30 sec on, 30 sec off). Lysate was clarified at 16,000xg, 15 min, 4 °C to remove insoluble debris. Protein concentration was measured by BCA assay (Pierce, 23227); thereafter, 50 µg protein per sample were used for all subsequent steps. Samples were reduced by DTT (final concentration 10 mM, 37 °C, 30 min, 700 RPM (Eppendorf Thermomixer R)), alkylated by IAA (final concentration 40 mM, room temperature, 45 min, protected from light), diluted 4x with 50 mM AmBic to reduce urea concentration below 2 M, and digested with trypsin (Pierce, 90305) in a 1:50 (w/w) ratio relative to sample protein (25 °C, overnight). Digestion was quenched by TFA (Sigma-Aldrich T6508; final concentration 1% v/v). Solid phase extraction was performed using SepPak C18 columns, 1 mL column volume, on a Zymo vacuum manifold. Cartridges were conditioned with 2x 1 mL 80% acetonitrile and 0.1% formic acid in LC-MS-grade Optima Water (Buffer B). Cartridges were equilibrated with 4x 1 mL 0.1% formic acid in Optima water (Buffer A). Cartridges were loaded with peptide samples, then washed with 4x 1 mL Buffer A and eluted into Buffer B by centrifuging the cartridges in a swinging bucket rotor, 300xg, 3 min. For each sample, the eluate was then transferred to a 1.5 mL Eppendorf tube and dried by vacuum centrifugation. Dried samples were stored at -80 °C until further processing.

PEG-independent purified viruses were resuspended in ∼250 uL of denaturing buffer and the proteins prepared for mass spectrometry as described above for infected cells.

### Mass spectrometry data acquisition

Dried peptide samples were resuspended in Buffer A (see above). Absorbance at A280 was measured for resuspended peptides to ensure that material was conserved during desalting. A280 values were not used to normalize the amount of material for LC-MS/MS injection, since the combined host-virus proteome of infected samples and the dramatically subset proteome of purified virions were unlikely to possess similar aromatic content. Instead, because a mass of 50 ug was digested for all samples, equal volumes were injected from each sample. Samples were separated by liquid chromatography using a Vanquish HPLC system (Thermo Fisher). Mobile phases consisted of Buffer A and Buffer B. For each run, 500 ng of peptides accumulated in a trap column (Acclaim Pepmap, 75 µm x 15 cm, 3 µm, 100 A) while the percentage of Buffer B increased linearly from 1% to 4% over 6 seconds, increased linearly from 4% to 9% over 8 minutes, and then held at 9% for 24 seconds. The trap column was then aligned with the resolving column (Acclaim Pepmap RSLC, 75 µm x 25 cm, 2 µm, 100 A). To elute peptides, the percentage of Buffer B was increased linearly from 9% to 23% over 30 minutes, then from 23% to 40% over 26 minutes. To wash the column, the percentage of Buffer B was increased linearly from 40% to 90% over 2.5 minutes, after which the percentage was held at 90% for 3 minutes. The chromatographic effluent was analyzed using a Thermo Orbitrap Ascend mass spectrometer in a data-independent manner. The MS1 spectra were collected using a scan range of 380-980 m/z, resolution of 120 K, maximum injection time of 251 ms, and AGC target of 1E6. MS2 spectra were acquired across a precursor scan range of 380-980 m/z in isolation windows of 12 m/z. Precursors were fragmented with an HCD collision energy of 30%. MS2 spectra were collected using a scan range of 150-2,000 m/z, resolution of 30 K, maximum injection time of 59 ms, and AGC target of 2.5E5. Raw proteomic data will be made available upon publication.

### Quantification of biofilm formation and cell attachment during infection with HFTV1

To quantify biofilm formation, an overnight *Hg* culture (<40 h) was diluted to OD_600_ ∼0.4 in 18% MGM and grown at 42 °C, 200 rpm until OD_600_ 1.0. The culture was infected with HFTV1 at MOI = 0, 1, 2, 10, then transferred to a 96-well plate with 200 μL per well. The plate was grown without shaking at 42 °C for 3 h. At 3h p.i., the supernatants were removed, and cells were fixed on the plate by drying at 60 °C for 1 h. Each well with fixed cells was stained with 200 μL of 0.05% crystal violet (Merck) for 30 min at ambient temperature. After staining, the crystal violet was removed, and the wells were washed twice with 300 μL of sterile ultrapure water. After drying, the crystal violet was solubilized by 150 μL of 33% acetic acid (Fisher Chemical). The quantity of biofilm was measured as absorbance at 595 nm using Multiscan FC (ThermoFisher Scientific) plate reader. For visual inspection of biofilm formation, *Hg* culture was grown as above and infected with HFTV1 at MOI = 0 and 10. The 5 mL cultures were transferred to glass test tubes and incubated at 42 °C for 3 h. At 3h p.i., the supernatants were removed, and the cells were fixed by drying at 60 °C for 1 h. Each tube was stained with 5 mL of 0.05% crystal violet (Merck) for 30 min at ambient temperature. After staining, the crystal violet was removed, and tubes were washed twice with 6 mL of sterile ultrapure water.

## QUANTIFICATION AND STATISTICAL ANALYSIS

### Differential expression analysis of Ribo-seq paired with RNA-seq

Differential expression (DE) analysis was performed using *DESeq2* [80] to identify genes that exhibited differential transcript abundance (dRNA) in RNA-seq or differential ribosome density (dRibo) in Ribo-seq. For both sequencing methods, read counts per feature (calculated using featureCounts [139]) were used to construct a *DESeqDataset* (DDS). A total of 18x RNA-seq libraries (3x biological replicates (BRs) at 0, 5, 15, 30, 60, and 180 min post-infection) and 16x Ribo-seq libraries (3x BRs at 0 min, 3x BRs at 5 min, 3x BRs at 15 min, 2x BRs at 30 min, 2x BRs at 60 min, and 3x BRs at 180 min) were included in this dataset. Since the *Hg* plasmid pHGLR3 was lost in one RNA-seq/Ribo-seq biological replicate, feature counts from pHGLR3 were isolated from all other genetic elements in both RNA-seq and Ribo-seq and used to construct separate DDSs. Due to the specific combination of Ribo-seq sample loss and plasmid loss, the 60 min time point is absent in the pHGLR3-specific DDSs. Processes applied to the non-pHGLR3 DDSs were similarly applied to pHGLR3-specific DDSs except where the 60 min time point was required. In the *DESeqDataset* metadata, the variables *batch* and *time* were assigned to each sample, with *time* treated as a nominal variable and *batch* included to account for batch effects. The *DESeqDataset* was constructed using *design = ∼ batch + time.* Genes were pre-filtered based on a count threshold of 50 and appearance in at least 2 samples. The data were rlog-transformed for quality assessment and various downstream visualizations.

Variance between samples was assessed by principal component analysis on rlog-transformed data. DE analysis was performed across all time points simultaneously using a likelihood ratio test (LRT) with *reduced = ∼ batch,* effectively asking, “Does variation in time account for variation in read counts?” We found this approach to be the most appropriate one for our time course data, since pairwise Wald hypothesis tests between all possible pairs of time points would entail an exorbitant number of tests and, by extension, a level of correction for false discovery rate that could inhibit the discovery of subtler expression changes. Resulting LRT adjusted p-values were corrected again post-hoc by the Benjamini-Hochberg procedure [153] within either RNA-seq or Ribo-seq to account for the number of tests conducted across all genetic elements (i.e., accounting for both pHGLR3 and non-pHGLR3 elements, since they were otherwise processed separately). These final significance values were then used to identify dRNA and dRibo genes based on a very stringent significance cutoff of adjusted p-value < 10e-5. Pairwise Wald hypothesis tests were also performed relative to 0 min post-infection for all other time points in both RNA-seq and Ribo-seq; however, these tests were simply used to extract log2 fold-change values for each gene and were not used for the identification of dRNA or dRibo genes.

### Cluster analysis of differentially expressed genes

Genes identified as dRNA or dRibo (see above) were clustered based on temporal expression pattern with *DP_GP_Cluster* [81]. *Rlog*-transformed, *sizeFactor-*normalized read counts from DESeq2 [80] were used as input; this data type has been previously shown to perform best in clustering steps compared to other forms of expression data [80]. Time in minutes was treated as “true”; in other words, time points were spaced according to the times at which samples were collected, rather than being spaced equidistant from each other. Clusters were formed over 1000 iterations. As a default processing step, gene expression patterns were mean-centered and scaled by *DP_GP_Cluster* so that similar patterns could be clustered with less interference from the magnitude of expression values. dRibo genes from pHGLR3 were processed separately from other genes for clustering due to the absence of a 60 min time point in the pHGLR3-specific data. In contrast, dRNA genes from all genetic elements were processed together for clustering, since data existed for at least two biological replicates containing all genetic elements at all time points. dRibo genes formed 27 clusters from non-pHGLR3 genes and 2 clusters from pHGLR3 genes. dRNA genes formed 23 clusters from all genetic elements. We then consolidated the gene clusters formed by *DP_GP_Cluster.* To that end, we calculated the mean trajectory or “silhouette” of each cluster from the genes comprising that cluster (based on *Rlog*-transformed, *sizeFactor-*normalized read counts (DESeq2)). The mean expression patterns of each cluster were then grouped in a second clustering step with *DP_GP_Cluster*, using a true time scale and 1000 iterations as before. This was a meta-clustering process of sorts, so we note that the output should not be interpreted as genuine clusters per se, at least in the context of the intended use for the *DP_GP_Cluster* program. For dRibo genes, the second clustering step produced 6 “cluster families” (CFs) that contained > 1 cluster. An additional CF was produced that contained only one cluster; this cluster was manually reassigned to one of the other CFs. Similarly, clusters of pHGL3 genes were manually reassigned to the 6x CFs. For dRNA genes, the second clustering step produced 6 CFs, accounting for all clusters and genetic elements. Qualitatively, dRibo CFs and dRNA CFs followed similar trajectories.

### Functional representation in cluster families

dRibo CFs were assessed for functional over- or under-representation of host functions based on functions from the arCOGs database [82], assigned via eggNOG v. 5.0 [154]. Representation of functions in CFs was relative to functions and their abundance in the *Hg* genome. First, the expected number of genes per functional category was calculated for each CF based on its size: expected number of genes per functional category = (total number of genes in CF)*(percent of genes for given functional category in the *Hg* genome). Next, for each functional category and each CF, the degree of representation was calculated by comparing the number of actual genes to their respective expected values: degree of representation = (actual number of genes for given functional category in CF)*(expected number of genes per functional category). Finally, the degree of representation was log-scaled for visualization, using a pseudocount to avoid the generation of undefined values, and reported as the "representation score" for simplicity: representation score = log2(degree of representation + 0.1). Significance of over- or under-representation of each function was tested within the *Hg* genes of each CF by binomial tests in R using *stats binom.test*. The number of successes, *x*, was the number of genes that appeared for a given functional category in the *Hg* genes of a CF; the number of trials, *n*, was the total number of *Hg* genes in a CF; and the probability of success, *p*, was the fraction of a given functional category in the entire *Hg* genome. Resulting p-values were corrected for false discovery rate by the Benjamini-Hochberg procedure [153] based on the total number of tests conducted for all functions and all CFs. The significance threshold for adjusted p-values was < 0.05. Here follows an illustration of the calculation described above using Cluster Family A and translation-related genes (arCOG J). There were 165 genes assigned to arCOG J in the *Hg* genome; this accounted for 4.09% of all protein-coding genes in the entire genome. Cluster Family A contained 783 *Hg* genes. Thus, the expected gene count for arCOG J in Cluster Family A was (783)*(0.0409) = 32.0. The actual number of arCOG J genes in Cluster Family A was 103, so degree of representation = 103/32 = 3.2; in other words, arCOG J was 3.2x more abundant in Cluster Family A than would be expected if 783 genes had been randomly selected from the *Hg* genome. The significance of this result was assessed using a binomial test, with *x* = 103, *n* = 783, and *p* = 0.0409, with a resulting p-value = 7.15E-25, or adjusted p-value = 5.00E-23 after BH-correction.

### Differential expression analysis for late-infection Ribo-seq

Ribo-seq libraries at 200 min post infection (2x biological replicates) and 150 min post-infection (3x biological replicates) were used to identify genes with differential ribosome density (dRibo) at late stages of infection (see “Ribo-seq sample collection” subheading “Ribo-seq at late-infection time point”). This analysis was performed with *DESeq2* [80]. The *DESeqDataset (DDS)* was constructed using read counts per feature from each library and *design = ∼ time*; all libraries used here were prepared in the same batch, so batch effects were not a concern and were excluded from the experimental design. Genes were pre-filtered based on a count threshold of 50 and appearance in at least 2 samples. The data were rlog-transformed for quality assessment and various downstream visualizations. Variance between samples was assessed by principal component analysis on rlog-transformed data. dRibo genes were identified using a pairwise Wald hypothesis test (contrasting 200 min vs. 150 min), with a significance threshold of 0.05 for the adjusted p-value and a magnitude threshold of 1.5 for log2 fold change. Functional analysis of *Hg* dRibo genes was based on annotations from the arCOGs database [82] via eggNOG v. 5.0 [154].

### Mass spectrometry data analysis

Fragpipe v. 24.0 [155] was used to analyze spectra of detected peptides. Match-between-runs ion false discovery rate was 1%. Spectra were searched against a combined HFTV1 and *Hg* proteome; the gp17 protein sequence was manually split in two according to the different transcript regions described in the main text, resulting in an N-terminal domain and a C-terminal domain treated as separate proteins. The proteome file will be made available upon publication. Spectral searches were filtered to peptide and protein FDRs of 1%. *In silico* digestion was performed by "stricttrypsin", allowing one missed cleavage. True precursor mass tolerance was +/-10 ppm, and MS2 fragmentation tolerance was +/-20 ppm. The spectral library produced by FragPipe was passed to DIA-NN (2.5.1 Academia) [156] for label-free quantification (LFQ), with a threshold for FDR of 1%. Protein-level LFQ DIA-NN output are available in **Table S11**.

Cumulative protein abundance derived from either the *Hg* or the HFTV1 proteome was calculated as the sum of all LFQ QuantUMS values [157] corresponding to proteins from *Hg* or HFTV1. To assess differential protein abundance of purified virions vs. infected cells 180 minutes post-infection, FragPipe-Analyst [158] was applied to the protein-level LFQ DIA-NN results. Proteins were considered differentially abundant based on a Benjamini-Hochberg adjusted p-value < 0.05 and log2 fold-change > |1.5|.

### qPCR fold-change calculation

Fold change was calculated by the ΔΔCt Livak method [159], using -HFTV1 as the control condition and *gyrB* as the reference gene.

